# The Integrity of the Speciation Core Complex is necessary for centromeric binding and reproductive isolation in *Drosophila*

**DOI:** 10.1101/2021.02.05.429932

**Authors:** Andrea Lukacs, Andreas W Thomae, Peter Krueger, Tamas Schauer, Anuroop V Venkatasubramani, Natalia Y Kochanova, Wasim Aftab, Rupam Choudhury, Ignasi Forne, Axel Imhof

## Abstract

Postzygotic isolation by genomic conflict is a major cause for the formation of species. Despite its importance, the molecular mechanisms that result in the lethality of interspecies hybrids are still largely unclear. The genus *Drosophila*, which contains over 1600 different species, is one of the best characterized model systems to study these questions. We showed in the past that the expression levels of the two hybrid incompatibility factors *Hmr* and *Lhr* diverged in the two closely related *Drosophila* species, *D. melanogaster* and *D. simulans*, resulting in an increased level of both proteins in interspecies hybrids. This overexpression leads to mitotic defects, a misregulation in the expression of transposable elements and a decreased fertility. In this work, we describe a distinct six subunit Speciation Core Complex (SCC) containing HMR and LHR and analyse the effect of *Hmr* mutations on complex function and integrity. Our experiments suggest that HMR acts as a bridging factor between centromeric chromatin and pericentromeric heterochromatin, which is required for both its physiological function and its ability to cause hybrid male lethality.

## INTRODUCTION

Eukaryotic genomes are constantly challenged by the integration of viral DNA or the amplification of transposable elements. As these challenges are often detrimental to the fitness of the organism, they frequently elicit adaptive compensatory changes in the genome. As a result of this process, the genomes as well as the coevolving compensatory factors can rapidly diverge between individuals within a population. Such divergences can result in severe incompatibilities eventually leading to the formation of two separate species (Presgraves, 2010; Sawamura, 2012).

Arguably the best characterized system for studying the genetics of reproductive isolation and hybrid incompatibilities is constituted by the two closely related *Drosophila* species *D. melanogaster* and *D. simulans* (*D. mel* and *D. sim*) (Sturtevant, 1920). One century of genetic studies has led to the identification of the three fast evolving genes that are critical for hybrid incompatibility: *Hmr* (Hybrid male rescue), *Lhr* (Lethal hybrid rescue) and *gfzf* (GST-containing FLYWCH zinc finger protein) (Hutter & Ashburner, 1987; Barbash & Ashburner, 2003; Watanabe, 1979; Brideau *et al*, 2006; Phadnis *et al*, 2015). The genetic interaction of these three genes results in the lethality of *D*.*mel*/*D*.*sim* hybrid males. Strikingly, all three genes are fast evolving and code for chromatin proteins suggesting that their fast evolution reflects adaptations to genomic alterations. While the molecular interaction between HMR and LHR is well established in pure species as well as in hybrids (Thomae *et al*, 2013; Satyaki *et al*, 2014), the molecular basis for their genetic interaction with GFZF is unclear. Interestingly, in interspecies hybrids or when HMR/LHR are overexpressed, HMR spreads to multiple novel bindings sites many of which have been previously characterized to also bind GFZF (Cooper *et al*, 2019).

In the nucleus of tissue culture cells and in imaginal discs, HMR and LHR form defined foci that are clustered around centromeres (Thomae *et al*, 2013; Kochanova *et al*, 2020; Blum *et al*, 2017). Super-resolution microscopy and chromatin immunoprecipitation revealed that HMR is often found at the border between centromeres and constitutive pericentromeric heterochromatin bound by HP1a (Kochanova *et al*, 2020; Gerland *et al*, 2017; Anselm *et al*, 2018). In addition to pericentromeric regions, HMR also binds along chromosome arms colocalizing with known *gypsy*-like insulator elements (Gerland *et al*, 2017). Depending on the tissue investigated, HMR shows slightly different binding patterns. It binds to telomeric regions of polytene chromosomes (Cooper *et al*, 2019; Thomae *et al*, 2013), colocalizes with HP1a in early *Drosophila* embryos (Satyaki *et al*, 2014) and near DAPI-bright heterochromatin in larval brain cells (Blum *et al*, 2017).

Flies carrying *Hmr* or *Lhr* loss of function alleles show an upregulation of transposable elements (TEs), defects in mitosis, and a reduction of female fertility in *D. mel*. Expression of transposable elements is increased particularly in ovarian tissue but also in cultured cells (Satyaki *et al*, 2014; Thomae *et al*, 2013). The mechanism that causes such a massive and widespread upregulation is not entirely clear as most of the TEs that respond to a reduced *Hmr* dosage are not bound by HMR under native conditions (Gerland *et al*, 2017). Due to the overexpression of the *HeT-A, TART* and *TAHRE* retrotransposons, *Hmr* mutants show a substantial increase in telomere length (Satyaki *et al*, 2014) and an increased number of anaphase bridges during mitosis presumably due to a failure of chromatid detachment during anaphase (Blum *et al*, 2017). The massive upregulation of transposable elements in ovaries is possibly also the cause of the substantially reduced fertility of *Hmr* and *Lhr* mutant female flies (Aruna *et al*, 2009).

Many of the phenotypes observed in cell lines lacking *Hmr* and *Lhr*, are mirrored by *Hmr* and *Lhr* overexpression, highlighting the importance of properly balanced *Hmr/Lhr* levels (Thomae *et al*, 2013). Hybrids show enhanced levels of both proteins relative to the pure species and consistently, are also characterized by loss of transposable elements silencing and cell cycle progression (Thomae *et al*, 2013; Kelleher *et al*, 2012; Satyaki *et al*, 2014). The latter is thought to be the cause for the failure of male hybrids to develop into adults, given the almost complete absence of imaginal discs (Blum *et al*, 2017; Gatti & Baker, 1989; Orr *et al*, 1997; Bolkan *et al*, 2007). In addition, hybrids and *Hmr/Lhr* overexpressing cells, display a widespread mis-localization of HMR at several euchromatic loci at chromosome arms including the previously unbound GFZF binding sites (Kochanova *et al*, 2020; Thomae *et al*, 2013; Cooper *et al*, 2019).

To better understand the deleterious effects observed in the presence of an excess of HMR, we decided to investigate the binding partners of HMR under native conditions and upon overexpression. Our results suggest that a defined Speciation Core Complex (SCC) of 6 subunits exists under native conditions and that interference with the formation of this complex results in its loss of function.

## MATERIALS AND METHODS

### Cell culture and induction

*Drosophila melanogaster* Schneider cell lines (SL2) were grown in Schneider’s medium (Gibco Life Technologies) supplemented with 10% fetal calf serum and antibiotics (penicillin 100 units/mL and streptomycin 100 µg/mL) at 26°C. Stable SL2 cells transfected with metallothionein promoter (pMT) driven *FLAG-HA-Hmr*^*+*^*/Myc-Lhr, FLAG-HA-Hmr*^*2*^*/Myc-Lhr, FLAG-HA-Hmr*^*dC*^*/Myc-Lhr, FLAG-HA-Boh1, FLAG-HA-Boh2* were generated as described in (Thomae *et al*, 2013). Cells carrying inducible transgenes were selected with 20 μg/mL Hygromycin B and induced for 18-24 h with CuSO_4_ before experiments. Cell lines carrying different *FLAG-HA-Hmr* transgenic alleles + *Myc-Lhr* as well as cells expressing *FLAG-HA-BOH1* were induced with 250 µM CuSO_4_. For *FLAG-HA-BOH2*, 500 µM CuSO_4_ was used. Wild type *FLAG-Hmr* SL2 cells used in Figure 2 and S3 were generated in (Gerland *et al*, 2017) by CRISPR-Cas9 mediated gene editing of the endogenous *Hmr* gene.

### Recombinant protein expression

*Spodoptera frugiperda* 21 (SF21) cells (Gibco Life Technologies) were used for baculovirus-driven recombinant co-expression of *HA-Hmr* and *His-Lhr*. Nuclei were extracted essentially as in (Dann *et al*, 2017) and HMR/LHR co-immunoprecipitation was performed using mouse anti-HA Agarose beads (1 μL packed beads/mL SF21 culture, Sigma-Aldrich A2095).

### Cloning

Cloning of *Hmr* and *Lhr* ORFs into the pMT-FLAG-HA plasmid was described in (Thomae et al., 2013). Restriction fragments containing *Hmr* and *Lhr* ORFs were sub-cloned into pFast Bac Dual (containing an N-terminal HA-tag) and pFast Bac HTb, respectively. The resulting plasmids were transformed into DH10Bac and recombined bacmids were isolated from clonal transformants. PCR verified recombinant bacmids were used for transfection of SF21 cells. *Boh1* and *Boh2* were PCR amplified from genomic DNA, cloned into pJet1.2 and verified by sequencing. ORFs were then sub-cloned into the pMT-FLAG-HA expression vector described in Thomae et al., 2013. Full cloning details, plasmids sequences and plasmids are available on request.

### Nuclear extraction for immunoprecipitation

Cells were harvested, centrifuged at 1200 x g and washed with cold PBS. Cell pellets were resuspended in hypotonic buffer (10 mM Hepes pH 7.6, 15 mM NaCl, 2 mM MgCl_2_, 0.1 mM EDTA, cocktail of protease inhibitors + 0.25 μg/mL MG132, 0.2 mM PMSF, 1 mM DTT) and incubated on ice for 20 min. Cells were incubated for another 5 min after addition of NP40 to a final concentration of 0.1% and then dounced with 20 strokes. 10% Hypertonic buffer (50 mM Hepes pH 7.6, 1 M NaCl, 30 mM MgCl_2_, 0.1 mM EDTA) was added to rescue isotonic conditions. Nuclei were centrifuged for 10 minutes at 1500 x g and the supernatant was discarded. Nuclei were washed once in isotonic buffer (25 mM Hepes pH 7.6, 150 mM NaCl, 12.5 mM MgCl_2_, 1 mM EGTA, 10% glycerol, cocktail of protease inhibitors + 0.25 μg/mL MG132, 0.2 mM PMSF, 1 mM DTT). After resuspension in the same buffer, nuclei were treated with benzonase (MERCK 1.01654.0001) and incubated for 30 minutes at 4°C on a rotating wheel. Soluble proteins were extracted by increasing the NaCl to 450 mM and incubated for 1 h at 4°C on a rotating wheel. Finally, the soluble material was separated from the insoluble chromatin pellet material by centrifugation for 30 min at 20000 x g and used for immunoprecipitations.

### Immunoprecipitation

Anti-FLAG immunoprecipitation was performed using 20 μL of packed agarose-conjugated mouse anti-FLAG antibody (M2 Affinity gel, A2220 Sigma-Aldrich) and were targeted either against the exogenously expressed transgenes (HMR^+^, HMR^dC^, HMR^2^) or an endogenously FLAG-tagged HMR (HMR). The other IPs were performed by coupling the specific antibodies to 30 µL of Protein A/G Sepharose beads. Each bait was targeted with at least one antibody (rat anti-LHR 12F4, mouse anti-HP1a C1A9, rabbit anti-NLP, anti-FLAG-M2 for FLAG-BOH1 and FLAG-BOH2), while HMR was targeted with three different antibodies (rat anti-HMR 2C10 and 12F1, anti-FLAG-M2 for FLAG-HMR). Rabbit anti-NLP and mouse anti-HP1a were directly incubated with the beads, while rat anti-HMR and anti-LHR were incubated with beads that were pre-coupled with 12 μL of a rabbit anti-rat bridging antibody (Dianova, 312-005-046). FLAG-IPs in non-FLAG containing nuclear extracts were used as mock controls for FLAG-IPs. For all other IPs, unspecific IgG coupled to Protein A/G Sepharose or Protein A/G Sepharose alone were used as mock controls.

The steps that follow were the same for all the immunoprecipitations and were all performed at 4°C. Antibody coupled beads were washed three times with IP buffer (25mM Hepes pH 7.6, 150 mM NaCl, 12.5 mM MgCl_2_, 10% Glycerol, 0.5 mM EGTA) prior to immunoprecipitation. Thawed nuclear extracts were centrifuged for 10 minutes at 20000 x g to remove precipitates and subsequently incubated with antibody-coupled beads in a total volume of 500-600 µL IP buffer complemented with a cocktail of protease inhibitors plus 0.25 μg/mL MG132, 0.2 mM PMSF, 1 mM DTT and end-over-end rotated for 2 h (anti-FLAG) or 4 h (other IPs) at 4°C. After incubation, the beads were centrifuged at 400 x g and washed 3 times in IP buffer complemented with inhibitors and 3 times with 50 mM NH_4_HCO_3_ before on beads digestion.

### Sample preparation for mass spectrometry

The pulled-down material was released from the beads by digesting for 30 minutes on a shaker (1400 rpm) at 25°C with trypsin at a concentration of 10 ng/μL in 100 µL of digestion buffer (1M Urea, 50 mM NH_4_HCO_3_). After centrifugation the peptide-containing supernatant was transferred to a new tube and two additional washes of the beads were performed with 50 μL of 50 mM NH_4_HCO_3_ to improve recovery. 100 mM DTT was added to the solution to reduce disulphide bonds and the samples were further digested overnight at 25°C while shaking at 500 rpm. The free sulfhydryl groups were then alkylated by adding iodoacetamide (12 mg/mL) and incubating 30 minutes in the dark at 25°C. Finally, the light-shield was removed and the samples were treated with 100 mM DTT and incubated for 10 minutes at 25°C. The digested peptide solution was then brought to a pH∼2 by adding 4 μL of trifluoroacetic acid (TFA) and stored at -20°C until desalting. Desalting was done by binding to C18 stage tips and eluting with elution solution (30% methanol, 40% acetonitrile, 0.1% formic acid). The peptide mixtures were dried and resuspended in 20 μL of formic acid 0.1% before injection.

### Sample analysis by mass spectrometry

Peptide mixtures (5 µL) were subjected to nanoRP-LC-MS/MS analysis on an Ultimate 3000 nano chromatography system coupled to a QExactive HF mass spectrometer (both Thermo Fisher Scientific). The samples were directly injected in 0.1% formic acid into the separating column (150 x 0.075 mm, in house packed with ReprosilAQ-C18, Dr. Maisch GmbH, 2.4 µm) at a flow rate of 300 nL/min. The peptides were separated by a linear gradient from 3% ACN to 40% ACN in 50 min. The outlet of the column served as electrospray ionization emitter to transfer the peptide ions directly into the mass spectrometer. The QExactive HF was operated in a Top10 duty cycle, detecting intact peptide ion in positive ion mode in the initial survey scan at 60,000 resolution and selecting up to 10 precursors per cycle for individual fragmentation analysis. Therefore, precursor ions with charge state between 2 and 5 were isolated in a 2 Da window and subjected to higher-energy collisional fragmentation in the HCD-Trap. After MS/MS acquisition precursors were excluded from MS/MS analysis for 20 seconds to reduce data redundancy. Siloxane signals were used for internal calibration of mass spectra.

### Proteomics Data analysis

For protein identification, the raw data were analyzed with the Andromeda algorithm of the MaxQuant package (v1.6.7.0) against the Flybase reference database (dmel-all-translation-r6.12.fasta) including reverse sequences and contaminants. Default settings were used except for: Variable modifications = Oxidation (M); Unique and razor, Min. peptides = 1; Match between windows = 0.8 min. Downstream analysis on the output proteinGroups.txt file were performed in R (v4.0.1). If not otherwise stated, plots were generated with ggplot2 package (v3.3.2). Data were filtered for Reverse, Potential.contaminant and Only.identified.by.site and iBAQ values were log_2_ transformed and imputed using the R package DEP (v1.10.0, impute function with following settings: fun= “man”, shift = 1.8, scale = 0.3). Except for Fig. 2 and 3, where data were bait normalized, median normalization was performed. Statistical tests were performed by fitting a linear model and applying empirical Bayes moderation using the limma package (v3.44.3). AP-MS for SCC identification (Fig. 1 and S1) were compared with a pool of all control samples (IgG and FLAG mock IPs). For Fig. 1C and Fig. S1E enriched proteins from AP-MS experiments from SCC components were first selected (cut off: log2FC > 2.5, p-adjusted < 0.05) and then intersection was quantified and plotted with UpsetR (v1.4.0). The Network graph in Fig. S1E was prepared with force directed layout in D3.js and R (Singh *et al*, 2020). The network graph in Fig. 2C was prepared using Cytoscape (3.4.0) with input from all AP-MS experiments and the String database (vs11.0) (Szklarczyk *et al*, 2017)

**Figure 1:**
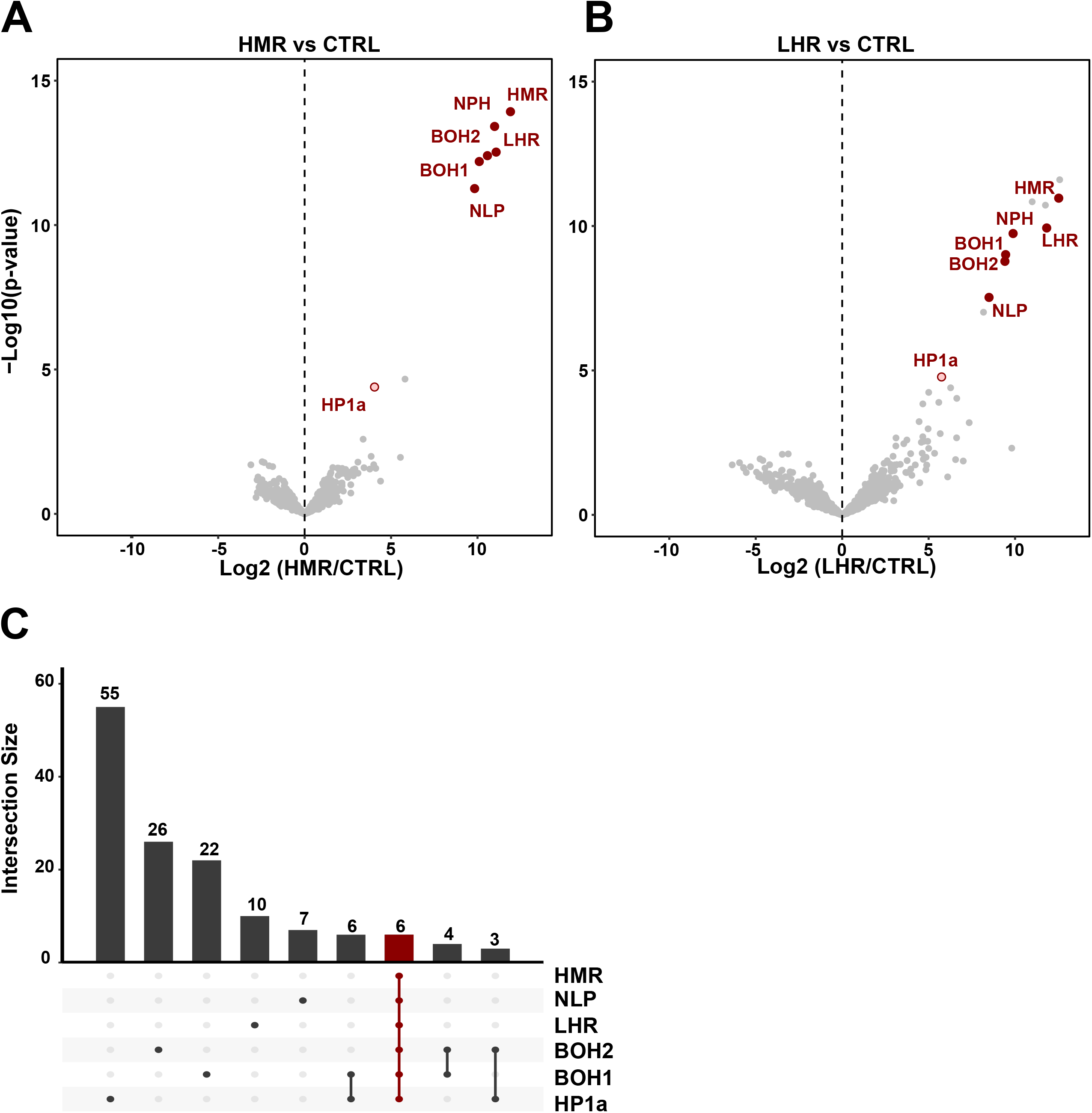
The hybrid incompatibility proteins HMR and LHR interact in the Speciation Core Complex (SCC). (**A**) and (**B**) Volcano plots highlighting reproducible interactors of HMR and LHR. Interactors consistently found in both HMR and LHR AP-MS are labeled in red and named the Speciation Core Complex (SCC). X-axis: log_2_ fold-change of factor enrichment in HMR (left, n = 8) or LHR (right, n = 4) IPs against mock purification (CTRL). Y-axis: significance of enrichment given as –log_10_ p-value calculated with a linear model. **(C)** The 6 SCC subunits are the only set of proteins shared in all HMR, LHR, NLP (n = 3), BOH1 (n = 4), BOH2 (n = 5) and HP1a (n = 4) AP-MS experiments (in red). Enriched proteins from each AP-MS experiment from SCC components were first selected (cut off: log2FC > 2.5, p-adjusted < 0.05) and then fed into UpsetR for intersection. Intersection plot showing the magnitude of the intersection (bars) among the different interactomes resulting from different IPs (rows). Lines connected dots define specific intersections between two or more interactomes. Unlabeled additional bait-specific interactors from **(A)** and **(B)** are available in Table S2 or in interactive plots at (<u>URL</u>). Additional volcano plots from SCC AP-MS are shown as supplementary Fig. S1.

**Figure 2:**
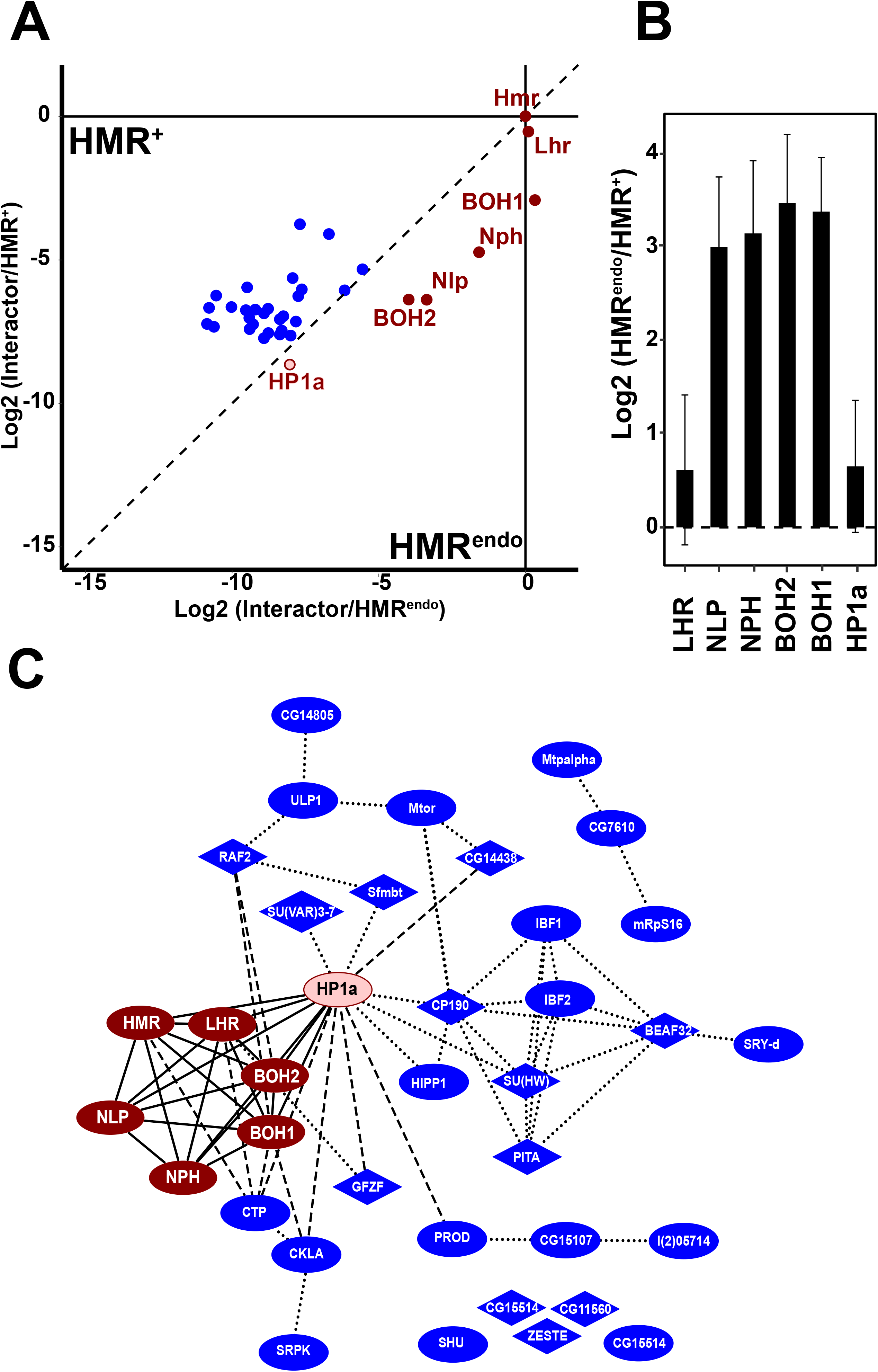
Overexpression leads excess of HMR to interact beyond the SCC. **(A**) Differential interaction proteome between endogenously FLAG tagged HMR (HMR^endo^, n = 4) and ectopically expressed HMR (HMR^+^, n = 9). Only proteins enriched in HMR^+^ or HMR^endo^ vs CTRL (p < 0.05) were considered. Components of the SCC and HP1a are shown in red, all other factors in blue. To display the differences within the HMR^endo^ and HMR^+^ interactomes, the enrichment of each putative interactor (Log2(iBAQ^HMR*^/IBAQ^control^)) was normalized to the enrichment of the HMR protein used as bait. The resulting values were then plotted against each other. Dots below the diagonal indicate a stronger enrichment in the HMR^endo^ pull down, the dots above the diagonal a stronger enrichment in the ectopically expressed HMR. (**B**) Differences of the ratio between HMR and members of the SCC and HP1a with and without ectopic expression of HMR. Plotted is the relative enrichment of each SCC member to the enrichment of the HMR used as bait (= the offset from the diagonal). Error bars reflect the standard error of the means (SEM). (**C**) Network diagram of all factors enriched in the AP-MS experiments of HMR^endo^ or HMR^+^ (p < 0.05). Red nodes represent SCC components and HP1a, blue nodes additional HMR binders. Nodes containing Zn-finger domains have a diamond shape. Solid edges connect the SCC and HP1a, dotted edges reflect protein-protein interactions predicted using the string database (Szklarczyk *et al*, 2017), dashed edges reflect protein-protein interaction identified by AP-MS experiments performed in this work (Fig. 1 and Table S3). In **(A)** and **(B)** proteins were labelled only if enriched in HMR^+^ or HMR^endo^ vs CTRL (p < 0.05).

**Figure 3:**
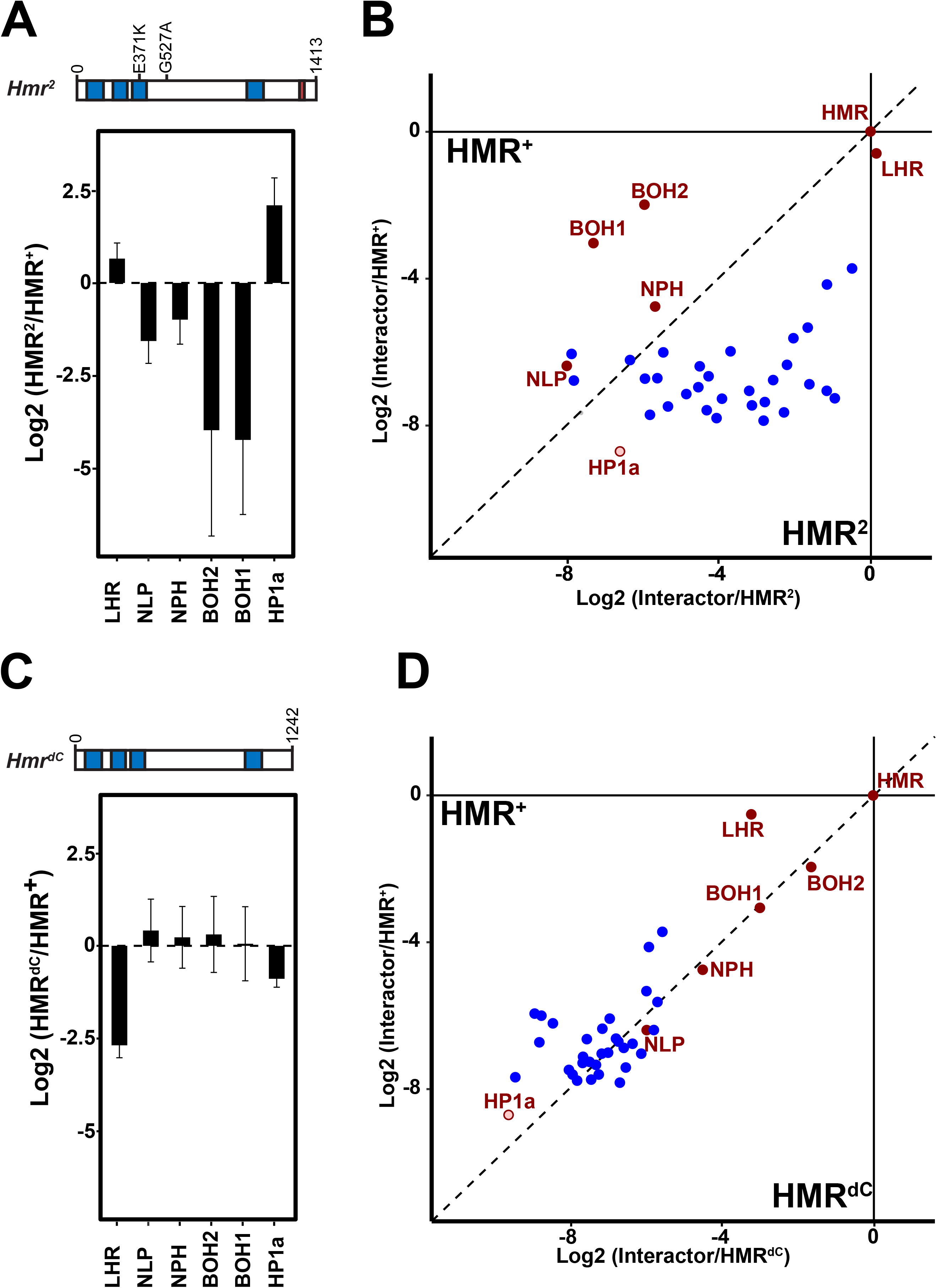
Two different *Hmr* mutations interfere differently with HMR interactome and SCC formation. Effect of *Hmr*^*2*^ **(A)** and *Hmr*^*dC*^ **(C)** mutations on the HMR interaction with the SCC components and HP1a. Y-axis represent the Log2 fold-change of HMR^2^/HMR^+^ and HMR^dC^/HMR^+^, respectively, calculated after normalization of each sample to the enrichment of the HMR protein used as bait. Error bars reflect the SEM (HMR^+^: n=9; HMR^dC^: n=10, HMR^2^: n=3). Differential interaction proteome between ectopically expressed wild type or mutated HMR (HMR^+^ versus HMR^2^ (**B**) or HMR^+^ versus HMR^dC^ (**D**)). Only proteins enriched in HMR^+^ or HMR^endo^ vs CTRL(p < 0.05) are shown. Components of the SCC are shown in red, all other factors in blue. To display the differences within each interactome, the enrichment of each putative interactor was normalized to the enrichment of the HMR protein used as bait. The resulting values were then plotted against each other. Dots above the diagonal indicate a stronger enrichment in the HMR^+^ pull down, dots below the diagonal a stronger enrichment in the HMR mutated alleles (HMR^2^ or HMR^dC^). Further details are available in Table S3.

### Western blot analysis

Samples were boiled 10 min at 96°C in Laemmli sample buffer, separated on SDS-PAGE gels, processed for western blot using standard protocols and detected using rat anti-HMR 2C10 (1:20), rat anti-LHR 12F4 (1:20), mouse anti-HP1a C1A9 (1:20), mouse anti-HA 12CA5 (1:20), mouse anti-FLAG SIGMA-M2 (1:1000), mouse anti-Lamin (1:1000) antibodies. Secondary antibodies included sheep anti-mouse (1:5000) (RRID: AB772210), goat anti-rat (1:5000) (RRID: AB772207), donkey anti-rabbit (1:5000) (RRID: AB772206) coupled to horseradish peroxidase. For proteins detection after IP, beads were boiled in Laemmli sample buffer after washing. For protein detection in ovaries, a short nuclear extraction was performed from 10 pairs of ovaries prior to boiling in sample buffer.

### ChIP-Seq

Chromatin immunoprecipitation was essentially performed as in (Gerland *et al*., 2017). For each anti-FLAG ChIP reaction, chromatin isolated from 1–2 × 10^6^ cells were incubated with 5 µg of mouse anti-FLAG (F1804, SIGMA-ALDRICH - RRID: AB262044) antibody pre-coupled to Protein A/G Sepharose. For ChIPs targeting total HMR, the same amount of chromatin was incubated with rat anti-HMR 2C10 antibody pre-coupled to Protein A/G Sepharose through a rabbit IgG anti-rat (Dianova, 312-005-046). Samples were sequenced (single-end, 50 bp) with the Illumina HiSeq2000. Sequencing reads were mapped to the *Drosophila* genome (version dm6) using bowtie2 (version 2.2.9) and filtered by mapping quality (-q 2) using samtools (version 1.3.1). Sequencing depth and input normalized coverages were generated by Homer (version 4.9). Enriched peaks were identified by Homer with the parameters -style factor -F 2 -size 200 for each replicate.

High confidence FLAG-HMR peaks (a pool of HMR^+^ and HMR^dC^) were called when a peak was present in at least half of the samples (5 out of 10). Coverages were centred at high confidence FLAG-HMR peaks in 4 kb windows and binned in 10 bp windows. The as-such generated matrices were z-score normalized by the global mean and standard deviation. HP1a-proximal peaks were defined as 10 percent of the peaks with highest average HP1a ChIP signal in 4 kb windows surrounding peaks. Composite plots and heatmaps indicate the average ChIP signal (z-score) across replicates. Heatmaps were grouped by HP1a class and sorted by the average ChIP signal in HMR native in a 400 bp central window. For statistical analysis, the average ChIP signal (z-score) was calculated in a 200 bp central window across peaks for each replicate. P-values were obtained by a linear mixed effect model (R packages: lme4 version 1.1-23 and lmerTest version3.1-2), in which average ChIP signal was included as outcome, genotype (*Hmr*^*+*^ or *Hmr*^*dC*^) and peak class (HP1a-proximal or non-HP1a-proximal) as fixed effects and sample ids as random intercept.

Chromosome-wide coverage plots were generated by averaging replicates, binning coverages in 50 kb windows and z-score normalizing by the global mean and standard deviation.

The percentage of peaks on chromosome 4 relative to the total number of peaks was calculated for each replicate. P-value was obtained by a linear model (R package: stats version 3.6.1), in which percentage was included as outcome and genotype (*Hmr*^*+*^ or *Hmr*^*dC*^) as independent variable.

### Immunofluorescent staining in SL2 cells

Immunofluorescent staining of SL2 cells was performed as described previously (Kochanova *et al*., 2020; Thomae *et al*., 2013). For Fig. 4A the following antibodies were used: mouse anti-HA 12CA5 1:1000 (Roche), rat anti-HMR 2C10 1:25 (Helmholtz Zentrum Munich), rabbit anti-dCENP-C 1:5000 (kind gift from C. Lehner) as a centromeric marker. For Figure S2A: mouse anti-HA (Invitrogen 2-2.2.14; 1:300), rabbit anti-CENP-A (Active Motif, 1:300), rat anti-HMR (2C10; 1:25); For Fig. S2B: rat anti-HA (Sigma Aldrich 3F10; 1:100), mouse anti-HP1a (DSHB C1A9, 1:100), rabbit anti-CENP-A (1:300). Fig. 4A: confocal microscopy z-scans were done on a Leica TCS SP5 (with 63x objective with 1.3 NA) with a step of 0.25 µM. Sum intensities projections were analyzed, and only cells with a minimum nucleoplasmic intensity of 70 a.u. on the anti-HMR channel were taken into account for analysis. Two different quantifications were performed. In one case cells were separated and counted based on the degree of co-localization between HMR and CENP-C: overlapping, partially overlapping or non-overlapping. In parallel, the number of CENP-C marked centromeric foci associated with HMR signal was measured. Both cells and centromeric foci were blind-counted, the experiment was repeated in 2 biological replicates and for each replicate at least two slides were measured (for each slide between 24 and 63 cells were quantified). Further details about stainings for Fig. S2 and microscopy are available as supplementary methods.

**Figure 4:**
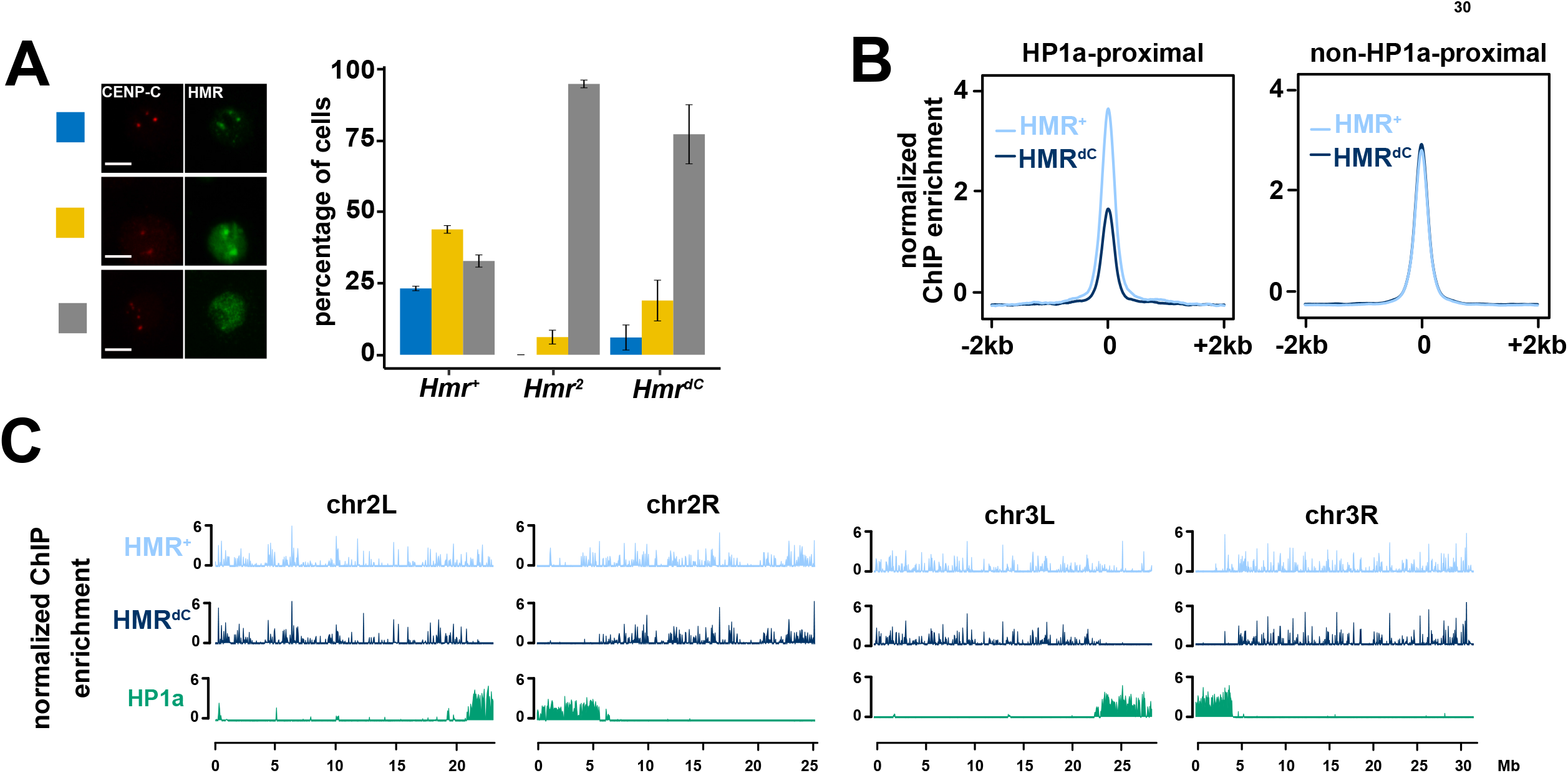
The HMR C-terminus is required for HMR localization in proximity to centromeres and HP1a-bound chromatin. (**A**) Ectopic HMR^dC^ fails to form bright (peri)centromeric foci in SL2 cells. Immunofluorescence images of cells expressing different *Hmr* transgenic alleles (HA-*Hmr*^*+*^, HA-*Hmr*^*dC*^ or HA-*Hmr*^*2*^) together with wild type LHR showing the co-staining of HA-HMR and CENP-C. Based on the overlap between HMR and CENP-C signals, cells were categorized in three groups (overlapping, partially-overlapping and non-overlapping) and the number of cells belonging to each group quantified. Error bars represent standard error of the means (n = 2). Size bars indicate 5 µm. (**B**) HP1a-proximal binding is specifically disrupted in HMR^dC^. Average plot of FLAG-HMR ChIP-seq profiles (z-score normalized) centered at high confidence FLAG-HMR peaks in 4 kb windows. HP1a-proximal (left) and non-HP1a-proximal (right) peaks are shown for HMR^+^ (light blue) and HMR^dC^ (dark blue). (**C**) HMR^dC^ genome-wide binding is impaired in proximity to centromeres and mostly unaffected at chromosome arms. Chromosome-wide FLAG-HMR ChIP-seq profiles (z-score normalized) for HMR^+^ (light blue), HMR^dC^ (dark blue) and HP1a (green). Chromosomes 2L, 2R, 3L and 3R are shown. Pericentromeric heterochromatin is marked by HP1a enriched territories distal (2L/3L) or proximal (2R/3R) to the respective x-axis. Plots in (**B**) and (**C**) represent an average of 5 biological replicates of FLAG-HMR ChIP-Seq in SL2 cells.

### Immunofluorescent staining in ovaries

Flies were grown at 25°C for 7-9 days and fed in yeast paste for at least 3 days prior to dissection. Ovaries were dissected in ice-cold PBS, then ovarioles were teased apart with forceps and moved to 1.5 mL tubes. PBS was removed and fixation solution (400 µL PBS Paraformaldehyde 2%, Triton 0.5 %, 600 µL Heptane) was added. Samples were incubated for 15 min at room temperature on a rotating wheel. Fixed ovaries were washed 3 times with 200 µL of PBS-T and then blocked for 30 min at room temperature on a rotating wheel (PBS-T, NDS 2%). After rinsing, 100 µL primary antibody solution (PBS-T, rat anti-HMR-2C10 1:20 mouse anti-HP1a-C1A9 1:10 and NDS 2%) were added and samples were incubated rotating overnight at 6°C. Primary antibody was washed three times with PBS-T. Secondary antibody solution was added (200 µL PBS-T + donkey anti-mouse Alexa 488 1:600, donkey anti-rat Cy3 1:300 and 2% NDS) and samples were rotated for 2 h at room temperature. Samples were washed three times with PBS-T and incubated for 10 min at room temperature with 200 µL of DAPI 0.002 mg/mL. Following this, they were washed once with PBS-T and once with PBS. Stained ovaries were finally mounted with one drop of vectashield in epoxy diagnostic slides (Thermo-Fisher Scientific, 3 wells 14 mm) and covered with high precision cover glasses. Further details about stainings for Fig. S7 and microscopy are available as supplementary methods.

### *Drosophila* husbandry and stocks

*Drosophila* stocks were reared on standard yeast glucose medium and raised at 25°C on a 12h/12h day/night cycle. Similar to (Thomae *et al*, 2013), the entire *D*.*mel* genomic region including the *melanogaster*-*Hmr* gene and parts of the flanking CG2124 and Rab9D genes (a 9538 bp fragment: X10,481,572-10,491,109), was cloned into a plasmid backbone containing a mini-white gene and a p-attB site. Plasmids for control *Hmr*^*+*^ stocks contained a wild type copy of *Hmr* while plasmids for test stocks contained either of the mutated versions *Hmr*^*dC*^ or *Hmr*^*2*^. *Hmr*^*dC*^ plasmids carry a point mutation with an A-T substitution (base 3667 of the CDS) that turns a Gly into a premature STOP codon and results in a C-terminally truncated protein product (last 171 aa missing). *Hmr*^*2*^ plasmids carry the two point mutations E371K and G527A (described in (Hutter *et al*, 1990; Aruna *et al*, 2009)). Identity of constructs was confirmed by sequencing. PhiC31 integrase-mediated transformations of the *D. melanogaster* line y1 w67c23; P{CaryP}attP2 (BL8622) were performed by BestGene Inc. resulting in the transgenic integration in attP2 docking site in chromosome 3 (3L:11070538). All rescue experiments were performed by crossing transgenes into *Df(1)Hmr-, ywv* background (Aruna *et al*, 2009)

### Crosses for generating *Hmr* genotypes for complementation tests in *D. melanogaster*

Males y w; *Hmr*^*\**^*/*TM6, Tb (*Hmr*^*\**^ = any *Hmr* transgenic allele, including *Hmr*^*+*^, *Hmr*^*dC*^, *Hmr*^*2*^) were crossed to *Df(1)Hmr-*females. F1 males *Df(1)Hmr-/Y; Hmr*^*\**^*/+* were backcrossed to the same females *Df(1)Hmr-* to generate females *Df(1)Hmr-/Df(1)Hmr-; Hmr*^*\**^*/+*. The latter females were crossed with their sibling males *Df(1)Hmr-/Y; Hmr*^*\**^*/+* to generate homozygotes for *Hmr*^*^ used for ovaries immunofluorescent HMR/HP1a stainings. To obtain isogenic flies for fertility and retrotransposon silencing complementation assays, heterozygous females *Df(1)Hmr-; Hmr*^*\**^*/+* were outcrossed to males *Df(1)Hmr-/Y* for 3-8 or 7-8 generations, respectively. Isogenized flies *Df(1)Hmr-; Hmr*^*\**^*/+* were crossed to *Df(1)Hmr-* and the resulting siblings were compared: control non-complemented individuals *Df(1)Hmr-* and test individuals *Df(1)Hmr-; Hmr*^*\**^*/+*.

### Crosses for generating *Hmr* genotypes for hybrid viability assays

Young *D*.*mel* females *Df(1)Hmr-*; *Hmr*^*\**^ were crossed at 25°C to 1-5 day old wild type *D*.*sim* males (C167.4 or *w*^*501*^). Control *D*.*mel* stocks *Df(1)Hmr-* were crossed in parallel to *D*.*sim* males (control cross: no lethality rescue). Crosses were transferred regularly to fresh medium. When larval tracks became visible, vials were transferred at 20°C to improve recovery of interspecific hybrids. Vials were kept until the last adults eclosed and number and genotype of hybrid offspring was scored. Rescue was measured by counting the number of viable transgene carrying males from the corresponding cross. In the rescue experiment, *Hmr*^*+*^ served as a positive (lethality rescue) and *Hmr*^*2*^ as a negative control (no lethality rescue). For statistical testing, Wilcoxon rank sum test (non-parametric) was used for pairwise comparisons with FDR correction for multiple testing using ggpubr package *(v0*.*4*.*0*, using compare_means function with following settings: formula = percent_males_offspring ∼ Hmr_allele, method = ‘wilcox.test’, p.adjust.method = ‘fdr’).

### Fertility assays

Three 1-3 days old *D. melanogaster* females were crossed for 2-3 days with six wild type males *D. melanogaster*. Flies were then transferred to fresh vials and again every 5 days for 3 times in total. Vials were scored 15-18 days after first eggs were laid, to make sure all adults were eclosed but no F2 was included. Vials in which one female or more than one male was missing were not scored. The whole assay was performed at 25°C. Tested females *Df(1)Hmr*-; *Hmr*^*\**^*/+* (*Hmr*^*^ = an *Hmr* transgenic allele) were always grown with and compared to their respective control siblings *Df(1)Hmr* ^-^;*+/+*, obtained from crosses between *Df(1)Hmr-* and *Df(1)Hmr*-; Hmr^*^/+. Rescue was measured as total offspring counted per female. In the rescue experiment, *Hmr*^*+*^ served as a positive control (fertility rescue) and Hmr^2^ as a negative control (no fertility rescue). For statistical testing, Wilcoxon rank sum test (non-parametric) was used for pairwise comparisons with FDR correction for multiple testing using ggpubr package *(v0*.*4*.*0*, using compare_means function with following settings: formula = offspring_per_mother ∼ Hmr_allele, group.by = ‘day’, method = ‘wilcox.test’, p.adjust.method = ‘fdr’).

### RNA extraction, cDNA synthesis and quantitative RT-PCR

2-10 pairs of ovaries were homogenized in Trizol (Thermo Fisher; cat. no. 15596026) and processed according to the manufacturer’s instructions. RNA concentration and A_260/280_ ratio were measured by NanoDrop. 1 μg of RNA was treated with DNase I recombinant, RNase-free (Roche; cat no.04716728001) followed by cDNA synthesis using SuperScript^™^III system (Invitrogen; cat. no: 18080051), both following the respective manufacturer’s protocols. For qPCR, equal volumes of cDNA for each sample were mixed with Fast SYBR^™^ Green Master Mix (Applied Biosystems^™^; cat. no: 4385610) and run in LightCycler® 480 Instrument II (Roche; cat. no: 05015243001) in a 384-well setup. Three technical replicates were used for each sample with 18s rRNA as the housekeeping gene. Annealing temperature for all the tested primers was 60°C and the list of primers used are given on request.

Plots for qPCR results were generated with R using ggplot2 package. For statistical testing Welch t-test was used with FDR correction for pairwise comparisons using rstatix package (v0.5.0).

### Data Sources

Published ChIP-seq data were obtained from GEO (HMR native and HP1a: GSE86106; CP190: GSE41354). New ChIP-seq datasets are available with the accession number: GSE163058. Proteomics datasets with the protein-protein interaction network of the SCC and the analysis of HMR overexpression and mutants are available with the accession numbers PXD023188 and PXD023193, respectively. Interactive network and volcano plots are available at Interactive Network and volcano plots from SCC purifications are entirely available at the following (URL) and in table S2.

## RESULTS

### Characterization of the Speciation Core Complex in *Drosophila melanogaster*

To identify the proteins interacting with the hybrid incompatibility (HI) proteins HMR and LHR under native conditions, we used specific monoclonal antibodies targeting HMR and LHR to perform affinity purification coupled with mass spectrometry (AP-MS) from nuclear extracts prepared from *D. mel* SL2 cells. For HMR, we additionally validated our results by performing AP-MS with a FLAG antibody in cells carrying an endogenously tagged HMR (HMR^endo^ (Gerland *et al*, 2017). These experiments revealed the existence of a set of four stable protein interactors shared between HMR (Fig. 1A) and LHR (Fig. 1B). Besides HMR and LHR, this six-subunit complex contains nucleoplasmin (NLP) and nucleophosmin (NPH) as well as two poorly characterized proteins, CG33213 and CG4788, which we named Buddy Of Hmr 1 (BOH1) and Buddy Of Hmr 2 (BOH2), respectively. AP-MS experiments for each of the individual subunits confirmed the existence of this defined complex (Figs. 1C, S1 A-E, Table S2) and the individual components also largely colocalize in SL2 cells (Fig. S2, (Anselm *et al*, 2018). As the two HI proteins HMR and LHR are involved in reproductive isolation and the formation of species, we termed the complex as Speciation Core Complex (SCC). HMR and LHR have been shown to interact and colocalize with the heterochromatin protein 1a (HP1a) (Greil *et al*, 2007; Brideau & Barbash, 2011; Satyaki *et al*, 2014; Thomae *et al*, 2013; Kochanova *et al*, 2020), which is consistent with our finding that both proteins as well as all other SCC subunits also interact with HP1a (Figs. 1C, S1A-E, Table S2). However, HP1a is only present in sub-stoichiometric amounts in the AP-MS purifications with antibodies against the SCC components, suggesting that it is not a stable component of the SCC. Individual AP-MS experiments also revealed proteins interacting exclusively with one or two components of the SCC (Fig. 1C and S1, Table S2), suggesting that the SCC components are also engaged in other complexes. In summary, our AP-MS results reveal the existence of a stable HMR/LHR containing protein complex (SCC) under physiological conditions. As several SCC subunits also contribute to other complexes, we wondered whether a surplus of HMR and LHR would affect complex composition.

### Overexpression of HMR and LHR results in a gain of novel protein-protein interactions

The importance of a physiological HMR and LHR dosage has been shown in non-physiological conditions like interspecies hybrids or cells where they are artificially co-overexpressed. The increased dosage of the two proteins results in their extensive genomic mislocalization (Thomae *et al*, 2013; Kochanova *et al*, 2020; Cooper *et al*, 2019). We therefore hypothesized that the overexpression of HMR and LHR results in a gain of novel interactions, which are potentially responsible for the novel binding pattern observed. Indeed, we confirmed 30 chromatin proteins that display a stronger interaction with HMR upon its overexpression (Thomae *et al*, 2013), Fig. 2A and S3, Table S2). These novel interactors include several proteins important for chromatin architecture such as the insulator proteins CP190, SU(HW), BEAF-32, IBF2 and HIPP1 or the mitotic chromosome condensation factor PROD. Thirteen of the novel interactors contain zinc finger DNA binding domains and three a MYB/SANT domain similar to HMR (Fig. 1C and Table S3). Intriguingly, one of the novel interactors is the product of the recently discovered missing third hybrid incompatibility gene, *gfzf*. Finding GFZF as an HMR interactor under expression conditions that resemble the situation in hybrids, provides a molecular explanation for its aberrant colocalisation with HMR both in hybrids and upon overexpression of HMR/LHR in tissue culture cells (Cooper *et al*, 2019).

We also found that the ratio between HMR/LHR and the other SCC components is lower in AP-MS experiments of ectopically expressed HMR/LHR (Fig. 2B,C). This suggests that the additional HMR and LHR molecules interact with other chromatin factors due to the limiting availability of the bona-fide SCC subunits. Notably, under these conditions the HMR/LHR/HP1a ratio is less affected by HMR/LHR overexpression than the interactions with NLP, NPH, BOH1 and BOH2.

Establishing to which extent these newly acquired interactors or the formation of a functional SCC contributes to HMR/LHR’s physiological function and it’s lethal function in male hybrids, would provide further mechanistic details. To this end we investigated the HMR interaction proteome upon ectopically expressing mutant HMR proteins.

### Two different *Hmr* mutations interfere with SCC formation and HMR localization

Most of the *Hmr* alleles that rescue hybrid male lethality are either null mutations or mutations that dramatically reduce the level of HMR (*Df(1)Hmr* ^*-*^, *Hmr*^*1*^, *Hmr*^*3*^, (Aruna *et al*, 2009; Hutter & Ashburner, 1987; Thomae *et al*, 2013)) and therefore don’t provide further mechanistic insights regarding Hmr’s role in hybrid incompatibility. The *Hmr*^*2*^ loss of function allele, however, just carries two point mutations: one within *Hmr*’s third of four MADF domains and one in an unstructured region of *Hmr*. This third MADF domain is unusual in that it is predicted to be negatively charged, which might mediate chromatin interactions rather than DNA binding (Aruna *et al*, 2009). Our previous experiments showed that HMR^2^ mislocalizes when expressed in SL2 cells (Thomae *et al*, 2013). To test whether these phenotypes can be explained by altered interaction partners, we expressed a FLAG-tagged HMR protein carrying the point mutations found in *Hmr*^*2*^ together with Myc-LHR in SL2 cells. A comparison of the interactome of ectopically expressed HMR^2^ and HMR wildtype FLAG fusion suggests that this mutation disrupts the interaction with the SCC components NLP, NPH, BOH1 and BOH2 while maintaining LHR and HP1a interactions (Fig. 3A). Interestingly most of the factors that are picked up by ectopic HMR appear to be more represented in HMR^2^ than in wildtype HMR purifications (Fig. 3B) suggesting that the interaction with novel interactors is not sufficient for HMR mediated lethality in hybrids.

Considering that the genetic interaction of *Hmr* and *Lhr* is critical for hybrid lethality (Brideau *et al*, 2006), and the previously established physical interaction between the two proteins, we asked whether interfering with their physical interaction would result in a loss of function. As the C-terminal BESS domain of HMR has been suggested to be responsible for the HMR/LHR interaction (Satyaki *et al*, 2014; Maheshwari *et al*, 2008; Brideau *et al*, 2006), we recombinantly expressed either wild type HMR or C-terminally truncated HMR (HMR^dC^) together with LHR using a baculovirus expression system. Supporting previous evidence of a direct HMR/LHR interaction mediated by HMR BESS domain, we could co-purify LHR only with full length HMR (Fig. S4A). Consistent with this observation *in vitro*, HMR^dC^ also shows a substantial reduction of interaction with LHR and HP1a when expressed in SL2 cells (Fig. 3C, Fig. S4B-C) while it still interacts with NLP, NPH, BOH1 and BOH2. Besides, a reduced interaction with LHR and HP1a, HMR^dC^ also interacts less efficiently with other heterochromatin components that are picked up upon HMR overexpression such as SU(VAR)3-7, HIPP1, PROD or CP190 (Fig. S4D-G, Table S3). These findings suggest that the deletion of the BESS domain does not lead to a complete disintegration of the SCC but specifically interferes with HMR’s binding to LHR, HP1a and other heterochromatin proteins. Therefore, the use of the *Hmr*^*dC*^ mutant allele allows us to selectively test for the functional importance of the HMR interaction with LHR and heterochromatin components and HMR’s potential ability to bridge different chromatin domains.

We next wondered whether the HMR C-terminal truncation and its concomitant loss of interaction with LHR and HP1a would influence its nuclear localization. A co-staining with antibodies against the exogenously expressed HMR and a centromeric marker (anti-CENP-C) revealed a rather diffuse nuclear localization of HMR^dC^, which is in sharp contrast to the full length HMR, which forms distinct bright (peri)centromeric foci (Fig. 4A, S6A) (Kochanova *et al*, 2020; Thomae *et al*, 2013).

To investigate the binding of ectopic HMR^dC^ to the genome, we performed a genome-wide ChIP-Seq profiling of HMR^dC^. HMR has been previously shown to have a bimodal binding pattern in SL2 cells (Gerland *et al*, 2017). One class of binding sites is found in proximity to HP1a containing heterochromatic regions whereas a second class is found along chromosome arms associated with *gypsy* insulators. Consistent with the failure to interact with HP1a, HMR^dC^ chromatin binding is specifically impaired at HP1a-dependent sites (Fig. 4B,C; Fig. S5) leading to a substantial reduction of HMR^dC^ binding in proximity to centromeres in both metacentric chromosomes 2 and 3 (Fig. 4B,C) as well as throughout the mostly heterochromatic chromosome 4 (Fig. S5C,D).

All together our results show that while the HMR C-terminus is required for HMR’s interaction with LHR and HP1a and for localization in close proximity to centromeres, it is dispensable for binding to the rest of the SCC and to genomic loci unrelated to HP1a. The *Hmr2* mutation in contrast does not affect HMR’s ability to interact with LHR/HP1a but weakens the interaction with the other SCC components. The fact that both mutations impair the centromere proximal binding suggests that complex integrity is necessary for HMR’s genomic localization. Next, we took advantage of these two mutants to investigate the importance of HMR’s interactions within the SCC for its function in flies.

### The HMR C-terminus is required for HMR physiological function in *D. melanogaster*

To investigate whether the C-terminus of HMR is required for HMR to fulfill its physiological function, we generated fly lines expressing full length HMR (HMR^+^) or mutant forms of it (HMR^dC^, HMR^2^). We crossed these alleles in a mutant background (*Df(1)Hmr-*, hereafter referred to as *Hmr*^*ko*^) and performed complementation assays to assess if the transgenic alleles are able to rescue *Hmr* wild type functions (Fig. 5A-B). All assays were done in ovaries, since HMR has been shown to be well expressed and important for fertility and retrotransposons silencing in this tissue. After verifying the expression of the *Hmr* transgenic alleles (Fig. S6D), we investigated their localization in follicle cells of sequentially developing egg chambers. Consistent with a tissue and potentially cell cycle dependent localization of HMR (Kochanova *et al*, 2020; Blum *et al*, 2017; Satyaki *et al*, 2014; Thomae *et al*, 2013), we find HMR colocalizing with CENP-C in early stage, mitotically cycling follicle cells of early-stage egg chambers (Fig. S7A). In post-mitotic and endoreplicating follicle cells of late-stage egg chambers lacking CENP-C, HMR localizes primarily to HP1a enriched domains (Fig. 5C). In contrast to what we observed in SL2 cells where the mutant proteins are expressed in the presence of an endogenous wild type copy, the centromeric localization of an HMR C-terminal mutant is only moderately affected in early stage follicle cells (Fig. S7B). However, in later stages HMR^dC^ does not associate with HP1a domains but rather shows a more diffuse nuclear staining, while the wild type transgenic HMR (HMR^+^) mirrors the localization of endogenous HMR (Fig. 5C).

**Figure 5:**
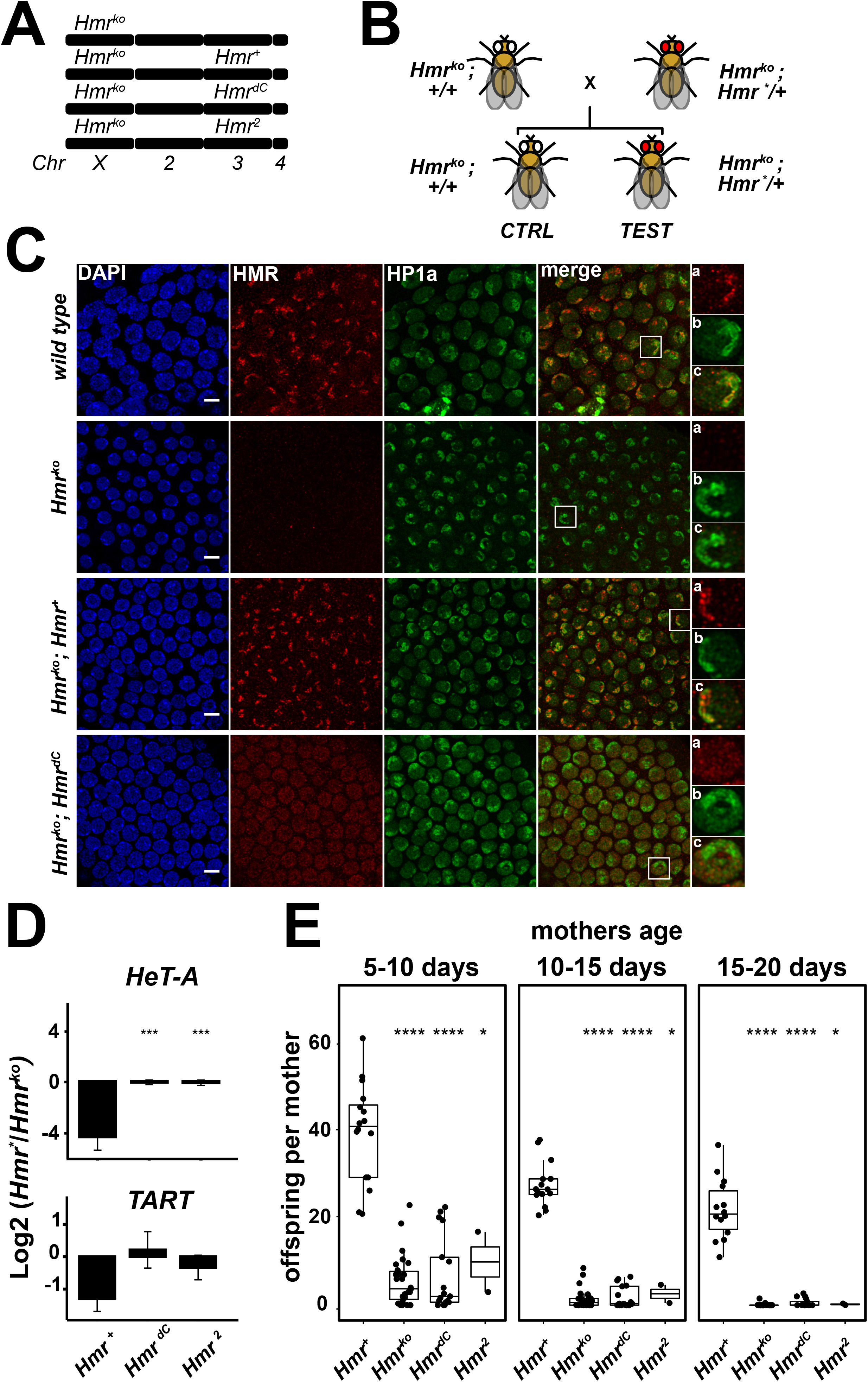
The HMR C-terminus is required for HMR physiological function in *D. melanogaster*. (**A**) Schematic representation of *Hmr* genotype in fly stocks used for complementation assays. *Hmr* null mutants (*Df(1)Hmr* ^*-*^, here referred to as *Hmr*^*ko*^) are complemented with wild type (*Hmr*^*+*^) or mutant (*Hmr*^*dC*^ and *Hmr*^*2*^) transgenic alleles inserted in the 3^rd^ chromosome. (**B**) Schematic representation of crosses performed for complementation assays. F1 siblings are compared: control animals (no *Hmr* transgene: *Df(1)Hmr* ^*-*^; +/+) and test animals (with *Hmr* transgene: *Df(1)Hmr* ^*-*^;*Hmr*^*^*/+*). *Hmr*^*\**^ refers to any transgene used in this work. (**C**) Truncation of the HMR C-terminus abrogates HP1a-proximal localization of HMR *in vivo*. Representative immunofluorescence images of stage 7 follicle cells in ovarioles from *Hmr* mutant *D. melanogaster* females complemented with either of the *Hmr* transgenic alleles. Insets represent zoom into one representative cell for anti-HMR (a), anti-HP1a (b) and merge of the two channels (c). Size bars indicate 5 µm. (**D**) Defective retrotransposons silencing in *Hmr*^*dC*^ and *Hmr*^*2*^. RT-qPCR analysis measuring mRNA abundance for the telomeric retrotransposons *HeT-A* and *TART* in female ovaries. Heterozygous genotypes (*Hmr*^*ko*^; *Hmr*^*\**^*/+)* were compared with the respective non-complemented siblings (*Hmr*^*ko*^; *+/+)*. Y-axis: log_2_ fold-change of the mean (complemented/non-complemented) after normalization to a housekeeping gene (18s-rRNA). Error bars represent standard error of the mean (n = 3). Welch t-test was used for pairwise comparisons with *Hmr*^*+*^ as a reference group and FDR for multiple testing adjustment (^*^ p ≤ 0.05, ^**^ p ≤ 0.01, ^***^ p ≤ 0.001, ^****^ p ≤ 0.0001). (**E**) HMR C-terminus is required for female fertility. Fertility defects are not complemented by *Hmr*^*dC*^: heterozygous females (*Hmr*^*ko*^; *Hmr*^*\**^*/+)* were compared with the respective non-complemented siblings (*Hmr*^*ko*^; *+/+)*. Number of adult offspring per female mother assessed in a time course (females aged 5-10, 10-15 and 15-20 days). Wilcoxon rank sum test was used for pairwise comparisons with *Hmr*^*+*^ as a reference group and FDR for multiple testing adjustment (^*^ p ≤ 0.05, ^**^ p ≤ 0.01, ^***^ p ≤ 0.001, ^****^ p ≤ 0.0001). For details about fertility assays refer to Table S4.

As the silencing of transposable elements (TE) has been previously shown to be impaired by *Hmr* loss of function mutations or knockdown (Satyaki *et al*, 2014; Thomae *et al*, 2013), we tested whether the *Hmr*^*dC*^ or the *Hmr*^*2*^ alleles were able to restore TE silencing (Fig. 5D). Whereas full length HMR was able to strongly repress all the TE studied, neither *Hmr*^*dC*^ nor *Hmr*^*2*^ were able to do so, showing expression levels comparable to *Hmr* deletion mutants. Since *Hmr* loss of function mutations have been shown to also cause a major reduction in female fertility (Aruna *et al*, 2009), we also tested *Hmr*^*dC*^ for the complementation of this phenotype. Similar to what we have observed for the TE silencing, both mutant alleles (*Hmr*^*dC*^ and *Hmr*^*2*^) were unable to rescue the fertility defect (Fig. 5E, Table S4). All together, these results show that the *Hmr* C-terminus is required for HMR localization and physiological function in *D. melanogaster* ovaries.

### The *Hmr* C-terminus is necessary for male hybrid lethality and reproductive isolation

To understand whether the toxic *Hmr* function in interspecies hybrids also requires its C-terminus, we performed a hybrid viability rescue assay (Fig. 6A). We therefore crossed *D. melanogaster* mothers carrying different *Hmr* alleles with wild type *D. simulans* stocks and counted the number of viable adult males in the offspring with respect to the total offspring (Fig. 6B, Table S5). In crosses with *Hmr* mutant mothers (*Df(1)Hmr* ^*-*^), hybrid male offspring counts are comparable to those of females. Introduction of a wild type *Hmr* transgene (*Hmr* ^*+*^), fully restores hybrid male lethality, while males carrying an *Hmr*^*dC*^ or an *Hmr*^*2*^ allele are viable. These results indicate that the *Hmr*^*dC*^, similarly to *Hmr*^*2*^, has lost its toxic function in hybrid males.

**Figure 6:**
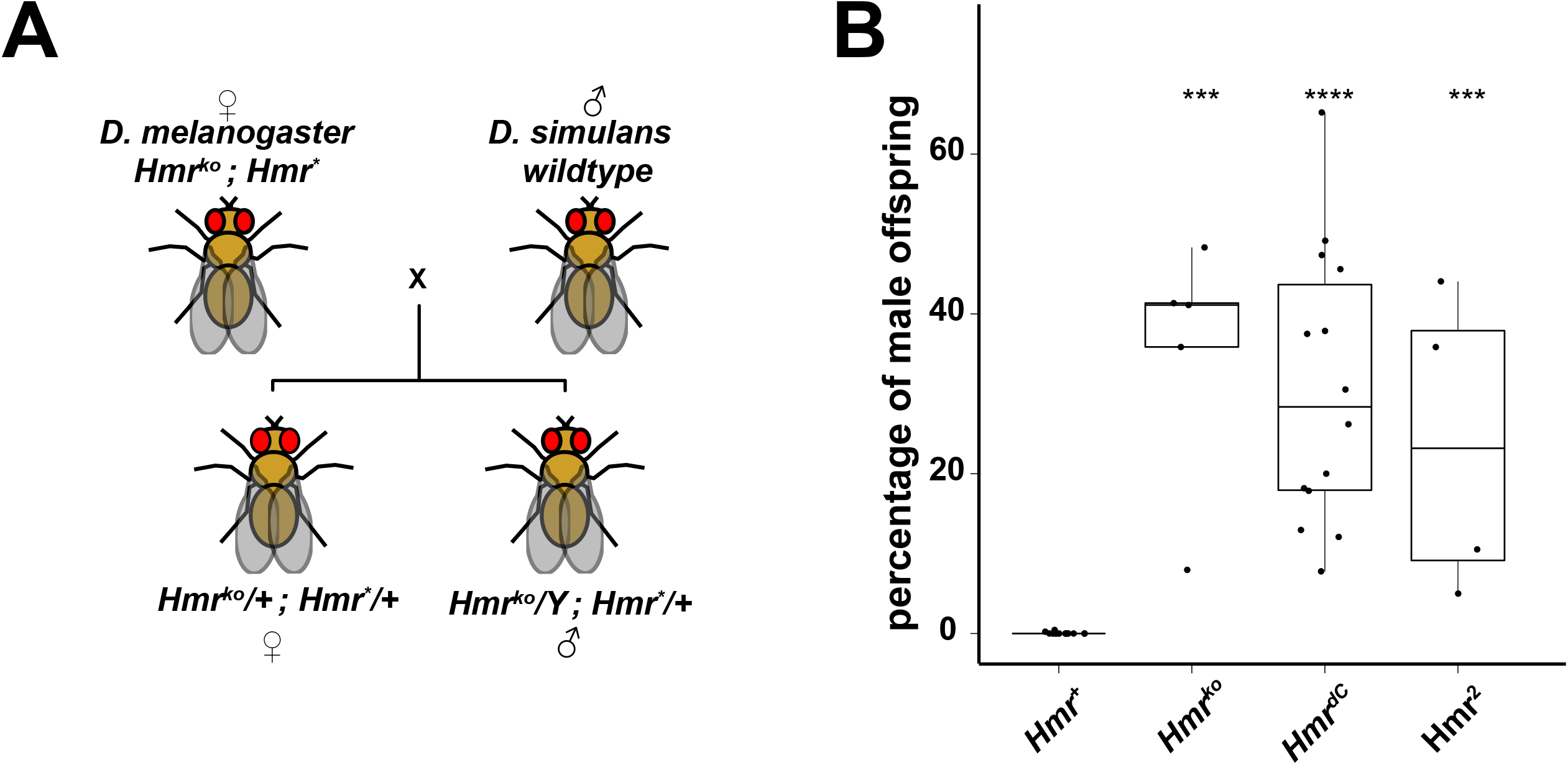
*Hmr* C-terminus is necessary for hybrid lethality. **(A)** Schematic representation of crosses performed for hybrid lethality rescue assays. Crosses between *D. melanogaster* mothers carrying *Hmr* transgenes in a *Df(1)Hmr* ^*-*^ background (*Hmr*^*ko*^; *Hmr*^*\**^) and wild type *D. simulans* males (C167.4 and *w*^*501*^). Hybrid male rescue is analyzed within the transgene-carrying offspring only. *Hmr*^*\**^ refers to any transgene used in this work. **(B)** HMR C-terminus is necessary for HMR lethal function in hybrid males. Suppression of hybrid males viability was measured as percentage of viable adult males in the total hybrid adult offspring. Crosses from non-complemented *Hmr*^*ko*^ mothers were used as control. Dots represent individual biological replicates. Wilcoxon rank sum test was used for pairwise comparisons with *Hmr*^*+*^ as a reference group and fdr for multiple testing adjustment (^*^ p ≤ 0.05, ^**^ p ≤ 0.01, ^***^ p ≤ 0.001, ^****^ p ≤ 0.0001). For details refer to Table S5.

## DISCUSSION

### HMR and LHR reside in a defined Speciation Core Complex (SCC)

HMR and LHR are two well-known *Drosophila* chromatin-binding factors whose overexpression results in male lethality in hybrid animals. When expressed at native levels in *D. melanogaster*, HMR and LHR form a distinct Speciation Core Complex (SCC) containing 6 subunits. This complex contains the two known histone chaperones NLP and NPH and the two yet uncharacterized proteins Buddy of HMR (BOH) 1 and 2 (CG33213 and CG4788) in addition to HMR and LHR. BOH1 is a putative transcription factor containing 4 Zn-finger DNA binding domains, which binds primarily to sites of constitutive (green) heterochromatin but also has a connection to components of (red) euchromatin (Bemmel *et al*, 2013). Many of these BOH1 binding sites are also bound by LHR (Filion *et al*, 2010) and HMR (Gerland *et al*, 2017), suggesting that BOH1 might play a role in recruiting the SCC to specific genomic loci (Liu *et al*, 2003; Kochanova *et al*, 2020; Blum *et al*, 2017) thereby bridging heterochromatic and euchromatic domains. While BOH1 contains 4 Zn-finger domains likely to bind DNA, no discernible domain can be identified in BOH2. Similar to HMR, LHR and BOH1 no ortholog of BOH2 can be identified outside the dipteran lineage and no molecular analysis of it has been done so far. Here, we show that BOH2 is a *bona fide* member of the SCC but also interacts with a set of other nuclear factors. *Nlp* and *Nph* are the two *Drosophila* paralogues of the nucleoplasmin family of histone chaperones, which are important for sperm decondensation upon fertilization (Emelyanov *et al*, 2014), chromosome pairing and centromere clustering (Joyce *et al*, 2012; Padeken *et al*, 2013). Both proteins depend on HMR to localize to the border between centromeric and pericentromeric chromatin (Anselm *et al*, 2018).

### Excess of HMR and LHR interact with novel chromatin factors

As hybrid animals suffer from increased levels of HMR and LHR, we performed AP-MS experiments of ectopically expressed HMR/LHR in SL2 cells in the presence of endogenous levels of HMR/LHR. This strategy allowed us to preferentially isolate proteins that interact with surplus molecules of HMR and LHR. Indeed, ectopically expressed HMR and LHR bind to a number of heterochromatic factors which are not detected under native conditions. Among many Zn-finger containing proteins that may explain the disperse localisation of overexpressed HMR on polytene chromosomes (Thomae *et al*, 2013) we observe a stable interaction of the extra HMR/LHR molecules with GFZF, the third factor required for male hybrid lethality (Phadnis *et al*, 2015). This finding is consistent with HMR and GFZF aberrantly colocalizing in interspecies hybrids and upon HMR/LHR overexpression (Cooper *et al*, 2019). Our interactome analysis with wild type and mutant HMR proteins suggests that HMR acts as a molecular bridge between different modules of the SCC.

### HMR contains two functionally important protein-protein interaction modules

Our proteomic analysis of *Hmr* mutants suggests that HMR’s N-terminal MADF3 domain, which is mutated in the *Hmr*^*2*^ allele, mediates the interaction with NLP, NPH, BOH1 and BOH2 while its C-terminus binds LHR, and through this interaction presumably recruits HP1a (Giot *et al*, 2003; Aruna *et al*, 2009; Thomae *et al*, 2013; Greil *et al*, 2007). Further genome-wide and cytological experiments with these *Hmr* alleles reveal that the integrity of all these interactions as well as a balanced expression of Hmr/Lhr is vital for proper targeting and the physiological functions of the SCC. Since the experiments in SL2 cells were done in the presence of endogenous (wild type) HMR, they reflect a competitive situation, which might be more sensitive in uncovering subtle differences between wild type and mutant HMR. Especially the centromere binding phenotype was more apparent in SL2 cells than in mitotically cycling follicle cells, in which the mutant *Hmr* allele was the sole source of HMR. The loss of heterochromatin association of HMR^dC^, however, was very apparent in follicle cells that have undergone a mitotic to endo-cycling transition. This finding argues that HMR interacts with HP1a through LHR to recruit the SCC to HP1a containing nuclear domains.

The functional assays in flies confirmed that both mutants are not able to rescue the HMR^ko^ phenotype, supporting the hypothesis that the integrity of the SCC complex is essential for its function. We therefore consider it unlikely that the *Hmr* mutant phenotype observed in *D*.*mel* (reduced female fertility, upregulation of TEs) is solely dependent on HMR’s ability to bind heterochromatin, since the mutant HMR^2^ protein is still able to interact with LHR and HP1a but no longer rescues these phenotypes. We therefore propose that HMR organizes the chromocenter by directly interacting with centromeric as well as heterochromatic factors and that both interactions are required for HMR to fulfil its function. In fact, defects in chromocenter bundling have been shown to result in micronuclei formation and loss of cellular viability in the imaginal discs and lymph glands (Jagannathan *et al*, 2019, 2018), a phenotype that is also observed in interspecies hybrids (Bolkan *et al*, 2007; Orr *et al*, 1997).

### HMR bridges heterochromatin and the centromere

Mutations impairing HMR’s bridging capacity between heterochromatin and the centromere not only fail to complement *Hmr* null phenotypes in pure species, but also do not cause lethality of male hybrids of *D*.*mel* and *D*.*sim*. This suggests that the simultaneous binding of HMR to heterochromatic and centromeric factors not only plays a role in its physiological function but also contributes to hybrid incompatibility. In addition, the novel finding that HMR/LHR overexpression leads to novel chromatin binders including GFZF (Thomae *et al*, 2013; Cooper *et al*, 2019) provides a possible molecular explanation for the requirement of all three proteins to mediate hybrid lethality (Phadnis *et al*, 2015). Based on our findings, we therefore propose that SCC integrity and the newly gained interactions play a key role in mediating hybrid lethality.

These newly gained protein-protein interactions that we observe in the presence of excessive HMR and LHR are probably the reason for an aberrant targeting of the SCC, a possible mislinkage of genomic loci and a failure to properly regulate the chromocenter in mitotically dividing cells. In combination these effects will eventually result in defects in cell cycle progression (Blum *et al*, 2017; Bolkan *et al*, 2007). As neither one of the described *Hmr* mutations is able to simultaneously bind heterochromatin and centromeric chromatin components, their expression does not result in hybrid lethality.

While we still do not understand the detailed molecular mechanism that mediate the physiological function of the newly defined SCC complex, our studies show that the complex has an important function in bringing two chromatin domains together. While the SCC might have evolved to perform this architectural function in *D*.*mel*, the postulated bridging function of HMR is clearly dispensable in *D*.*sim*. In hybrids where increased levels of HMR and LHR confronted with two different genomes that have independently evolved, they are involved in a number of novel aberrant molecular interactions that result in lethality.

The isolation of a defined complex involving the hybrid incompatibility proteins HMR and LHR will enable us to perform a more detailed molecular analysis of its function and sets the ground for future comparative studies on the divergent evolution of its components within species and on their lethal interactions in hybrids.

## DATA AVAILABILITY

ChIP-seq datasets are available with the accession number: GSE163058. Proteomics datasets with the protein-protein interaction network of the SCC and the analysis of HMR overexpression and mutants are available with the accession numbers PXD023188 and PXD023193, respectively. Interactive Network and volcano plots from SCC purifications are entirely available at the following https://wasim-aftab.shinyapps.io/SccNet-AL/.

## SUPPLEMENTARY DATA

Supplementary Data are available online.

## ACKNOWLEDGEMENTS

We thank Patrick Heun, Christian Lehner and Harald Saumweber for the Nlp, CENP-A, CENP-C and Lamin antibodies respectively. We also thank Daniel Barbash for the Df(1)Hmr-fly strain. We furthermore would like to thank Catherine Regnard, Sandro Baldi and Alessandro Scacchetti for critical comments on the manuscript. Andreas Schmidt and Marc Wirth for advice on proteomic analysis and for mass-spectrometry analysis. Irene Vetter and Silke Krause for cloning of expression constructs. Thomas Gerland, Raffaella Villa and Alessandro Scacchetti for advice on ChIP-seq, Angelika Zabel on ChIP-seq library preparation and Kenneth Boerner for advice on ovary staining. We also would like to thank Sophia Groh for graphical aid and Elisabeth Schröder-Reiter and Markus Hohle for constant support. We also would like to thank the entire Imhof group, Peter Becker and the Becker department for helpful discussions. In addition, we would also like to thank Stefan Krebs and Helmut Blum from LAFUGA facility for sequencing.

## SUPPLEMENTARY FIGURES

**Figure S1:**
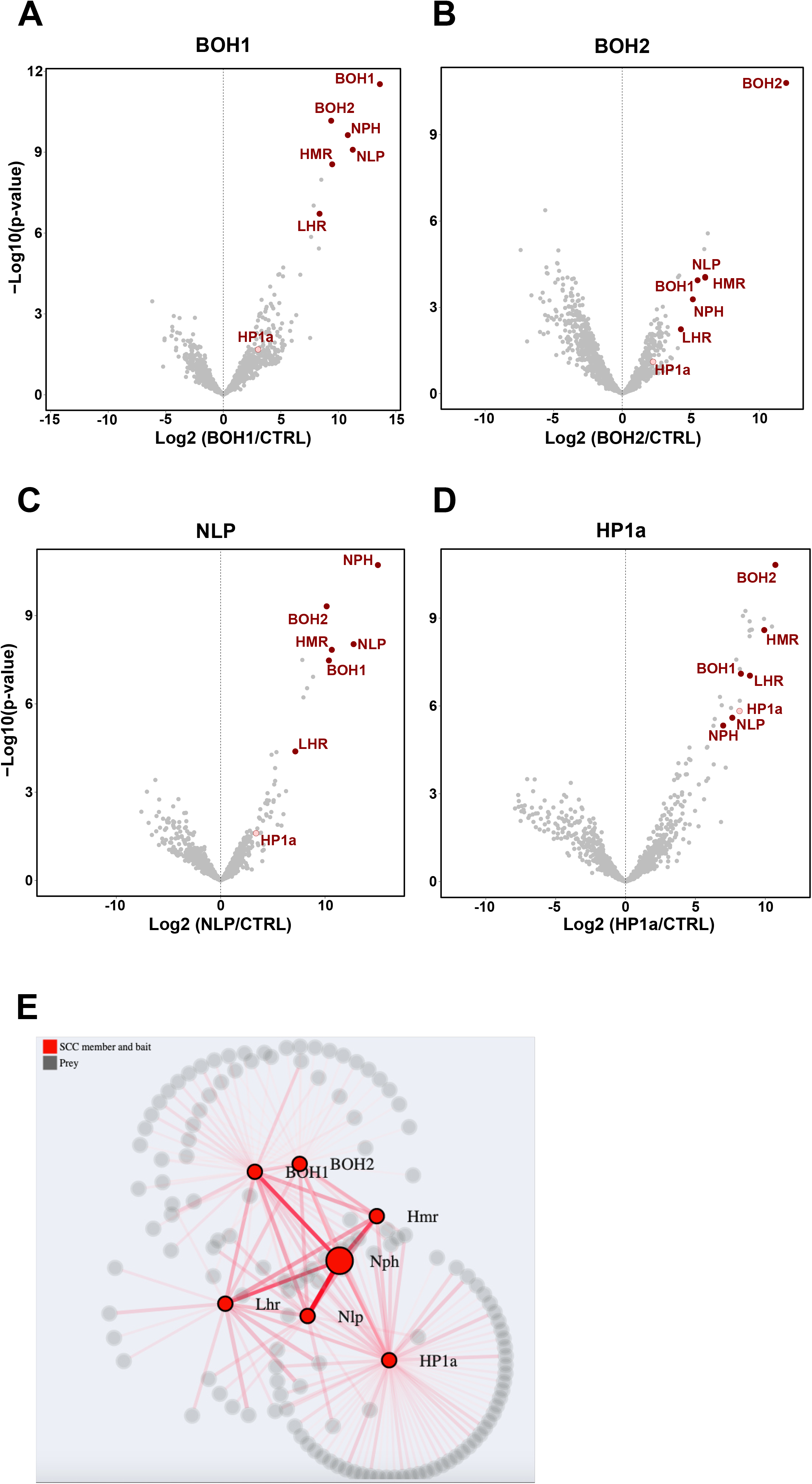
Interactomes of HMR/LHR interactors provide evidence for the existence of a Speciation Core Complex (SCC) (related to Fig. 1). **(A**) - (**D**) SCC proteins are consistently enriched in reciprocal IPs. Volcano plots showing the interactome resulting from the AP-MS of the SCC proteins BOH1 (n = 4), BOH2 (n = 5), NLP (n = 3) and HP1a (n = 4) against a “mock” purification. The SCC components are labeled in red. X-axis: log_2_ fold-change of factor enrichment in IPs against mock purification (CTRL). Y-axis: significance of enrichment given as –log_10_ p-value calculated with a linear model. A list of the unlabeled additional bait-specific interactors is provided in Table S2. (**E**) Network plot showing a highly connected SCC surrounded by subunit-specific interactors. Enriched proteins from each AP-MS experiment from SCC components were first selected (cut off: log2FC > 2.5, p-adjusted < 0.05) and integrated in an <u>interaction network</u> drawn with force directed layout in D3.js and R. Nodes represent proteins significantly interacting with at least one of the SCC components. Edges represent physical connections experimentally detected in this work. SCC components (and baits) are labelled in red. Interactive volcano plots and interaction network are available at the following (<u>URL).</u>

**Figure S2:**
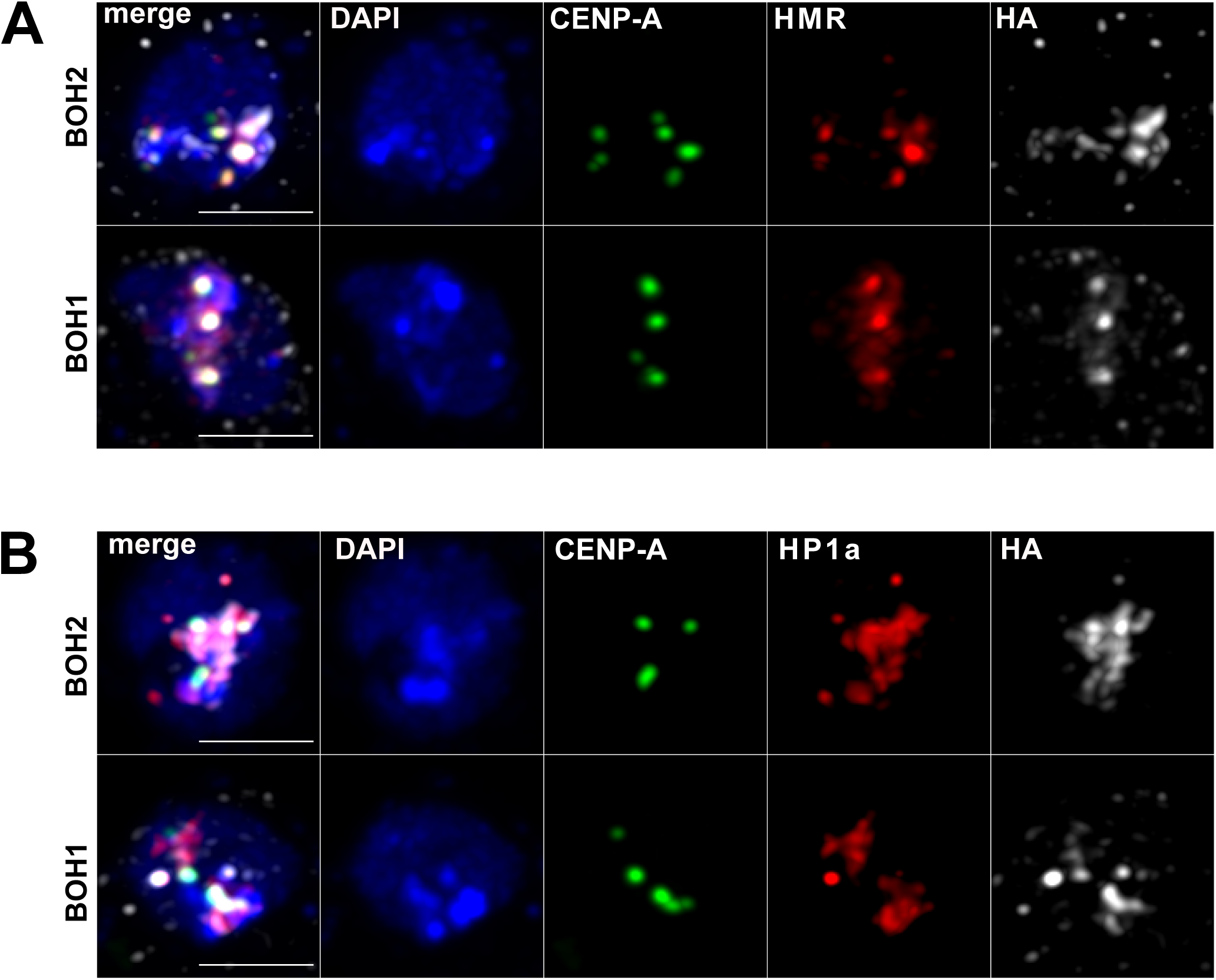
The SCC components BOH1 and BOH2 largely colocalize with HMR in proximity to centromeres and HP1a domains (related to Fig. 1). **(A)** BOH1 and BOH2 colocalize with HMR and CID in SL2 cells. Immunofluorescence images of cells expressing FLAG-HA-BOH2 (upper panel) or FLAG-HA-BOH1 (lower panel) showing the co-staining of HA-BOH2 or HA-BOH1, respectively, with CID (centromeres) and HMR. **(B)** BOH1 and BOH2 colocalize with HP1a and CID in SL2 cells. Immunofluorescence images of cells expressing FLAG-HA-BOH2 (upper panel) or FLAG-HA-BOH1 (lower panel) showing the co-staining of HA-BOH2 or HA-BOH1, respectively, with CID (centromeres) and HP1a (pericentromeric chromatin). For **(A)** and **(B)** size bar indicates 3 µm, DAPI staining indicate nuclei.

**Figure. S3:**
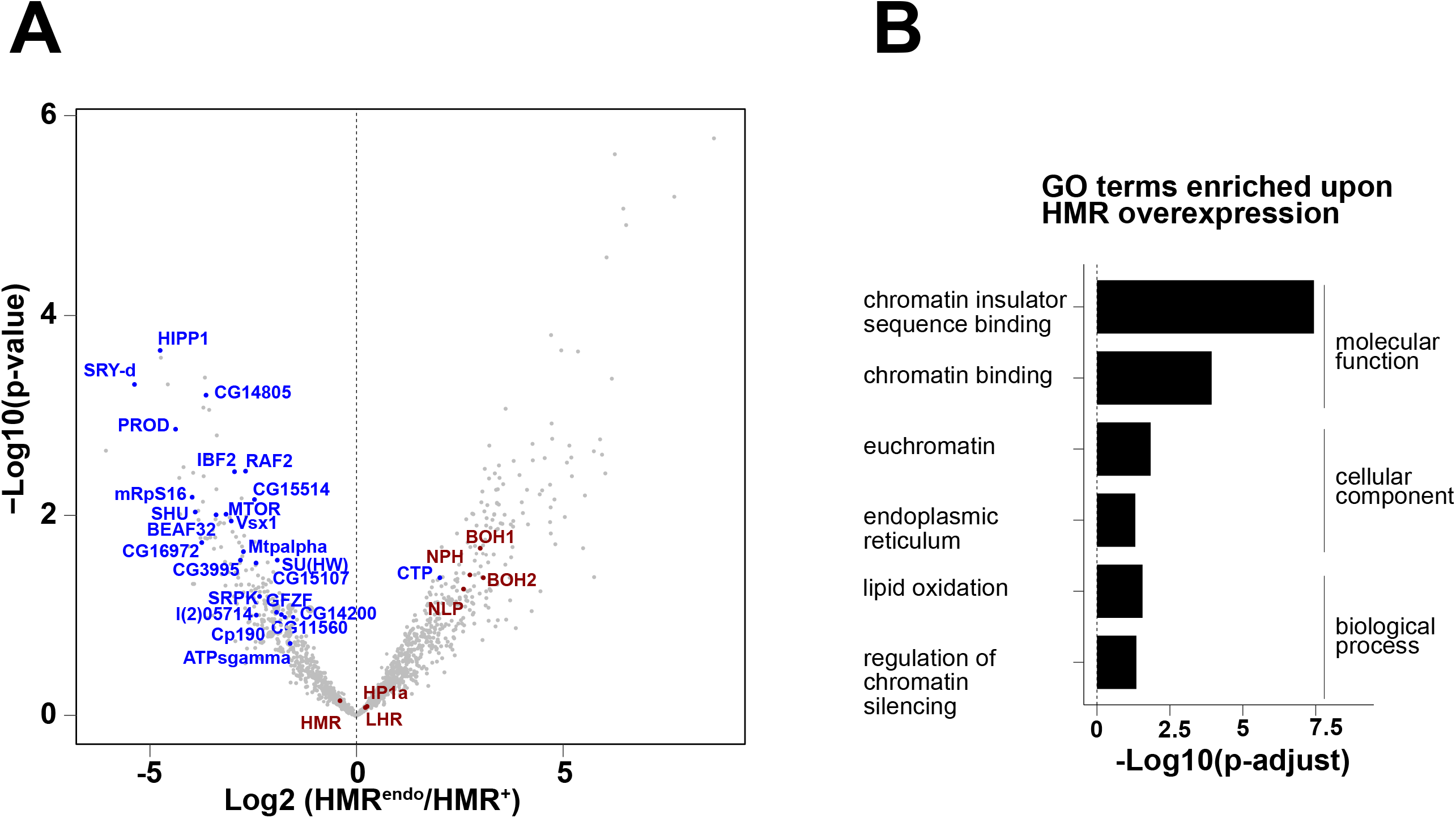
Excess of HMR interacts beyond the SCC with chromatin architecture proteins (related to Fig. 2). **(A)** Volcano plot highlighting novel interactions gained by HMR upon overexpression. X-axis: log_2_ fold-change of FLAG-HMR^endo^ IPs (right side of the plot) vs FLAG-HMR^+^ IPs (left side of the plot). Y-axis: significance of enrichment given as –log_10_ p-value calculated with a linear model. SCC subunits are labelled in red, novel factors enriched upon HMR overexpression are labelled in blue. Unlabeled additional bait-specific interactors are listed in Table S3. **(B)** GO terms enriched upon overexpression of HMR. In **(A)** and **(B)** proteins were labelled or considered for GO search only if enriched in HMR^+^ or HMR^endo^ vs CTRL (p < 0.05) and differentially enriched between HMR^+^ and HMR^endo^ (log_2_ fold-change (HMR^endo^/HMR+) < 1.5).

**Figure S4:**
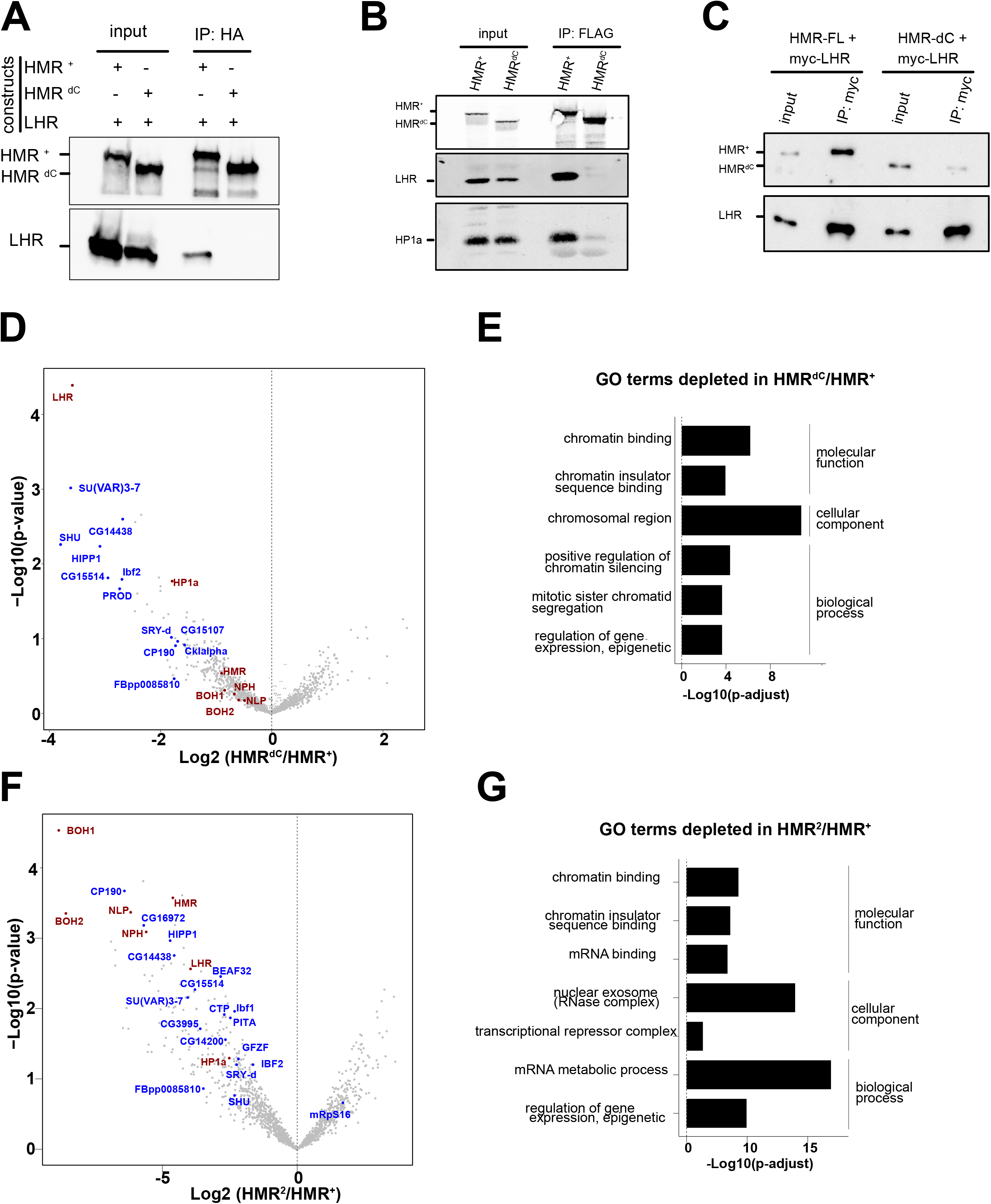
Two different *Hmr* mutations interfere differently with HMR interactome and SCC formation (related to Fig. 3) **(A)** Recombinantly co-expressed HMR^dC^ and LHR do not interact. Western blot showing anti-HA immunoprecipitation in nuclear extracts from Sf21 insect cells transfected with HA-HMR and His-LHR. IP performed with anti-HA antibody and western blot probed with anti-HMR or anti-LHR antibodies. **(B)** HMR C-terminus is required for HMR interaction with LHR and HP1a in *D. mel* SL2 cells. Western blot showing HMR immunoprecipitation in SL2 cells stably transfected with either full length HMR or a C-terminally truncated HMR^dC^ along with Myc-LHR. IP performed with anti-FLAG antibody targeting FLAG-HMR and western blot probed with anti-HA (HMR), anti-Myc (LHR) and anti-HP1a. **(C)** Western blot showing LHR immunoprecipitation in SL2 cells stably transfected with either full length HMR or a C-terminally truncated HMR^dC^ along with Myc-LHR. IP performed with anti-Myc antibody targeting Myc-LHR and western blot probed with anti-FLAG (HMR) and anti-LHR. **(D)** Volcano plot highlighting interactions depleted in HMR^dC^. X-axis: log_2_ fold-change of FLAG-HMR^dC^ IPs (right side of the plot) vs FLAG-HMR^+^ IPs (left side of the plot). Y-axis: significance of enrichment given as –log_10_ p-value calculated with a linear model. SCC subunits are labelled in red. In blue are factors depleted upon HMR^dC^ mutation (among the endogenous or overexpression-induced interactions of HMR). Unlabeled additional bait-specific interactors are listed in Table S3. **(E)** GO terms depleted upon HMR^dC^ mutation. **(F)** Volcano plot highlighting interactions depleted in HMR^2^. X-axis: log_2_ fold-change of FLAG-HMR^2^ IPs (right side of the plot) vs FLAG-HMR^+^ IPs (left side of the plot). Y-axis: significance of enrichment given as –log_10_ p-value calculated with a linear model. SCC subunits are labelled in red. In blue are factors depleted upon HMR^2^ mutation (among the endogenous or overexpression-induced interactions of HMR). Unlabeled additional bait-specific interactors are listed in Table S3. **(G)** GO terms depleted upon HMR^2^ mutation. In **(D) and (G)** proteins labelled or used for GO search include endogenous or overexpression-induced interactions of HMR (i.e. enriched in HMR^+^ or HMR vs CTRL with p < 0.05) and differentially enriched between HMR^+^ and the HMR mutant analyzed (log_2_ fold-change (HMR^*^/HMR+) < 1.5).

**Figure S5:**
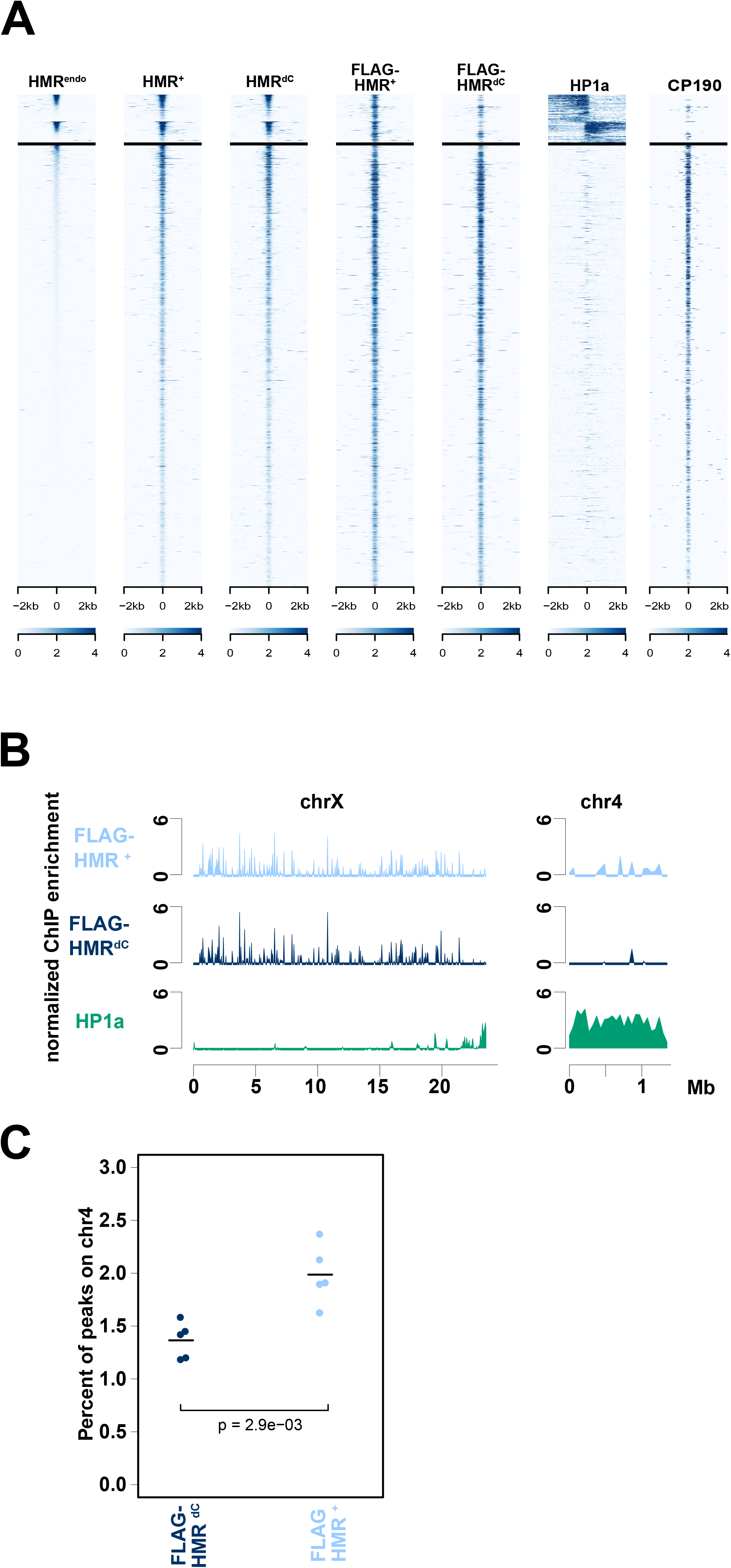
The HMR C-terminus is required for HMR localization in proximity to centromeres and HP1a-bound chromatin (related to Fig. 4) **(A)** Heatmaps of ChIP-seq profiles (z-score normalized) centred at high confidence FLAG-HMR peaks in 4 kb windows. Peaks are grouped by HP1a class and sorted by the ChIP signal in native HMR ChIP. From left to right, anti-HMR ChIP in untransfected cells, anti-HMR ChIP in cells transfected with FLAG-*Hmr*^+^ and FLAG-*Hmr*^*dC*^, anti-FLAG ChIP of cells transfected with FLAG-*Hmr*^+^ or FLAG-*Hmr*^*dC*^, and anti-HP1a and anti-CP190. The latter two are representative of the two classes of HMR peaks: HP1a-proximal and non-HP1a-proximal. **(B)** Chromosome-wide FLAG-HMR ChIP-seq profiles (z-score normalized) for *Hmr*^+^ (light blue), *Hmr*^*dC*^ (dark blue) and HP1a (green). Chromosomes X and 4 are shown.**(C)** *Hmr*^*dC*^ is depleted at heterochromatin rich chromosome 4. Percentage of FLAG-HMR ChIP-seq peaks located on chromosome 4 for each replicate (n=5). *Hmr*^*+*^ (light blue) and *Hmr*^*dC*^ (dark blue) are shown. P-values are obtained by a linear model. FLAG-HMR plots represent an average of 5 biological replicates.

**Figure S6:**
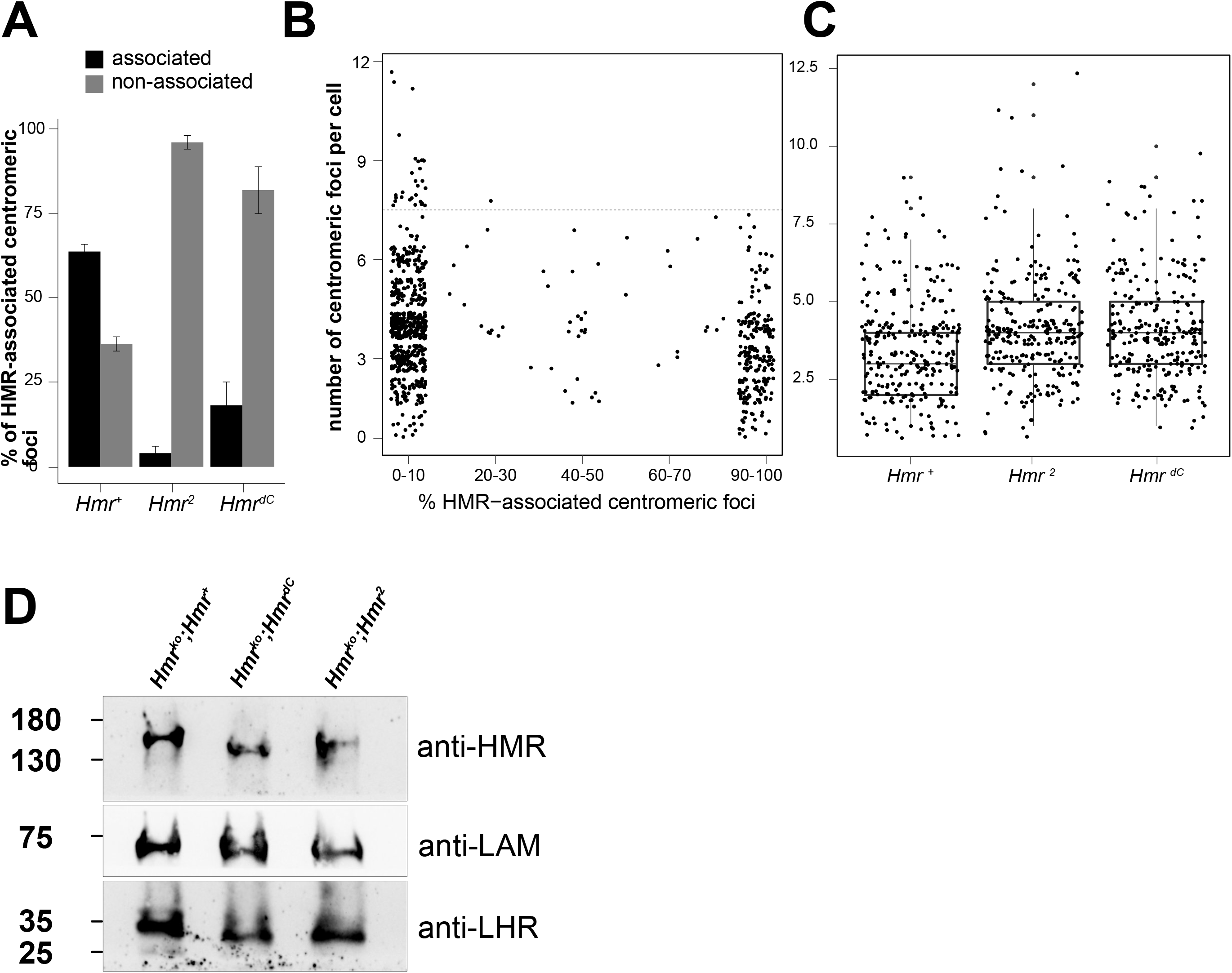
The HMR C-terminus is required for HMR localization in proximity to centromeres and HP1a-bound chromatin (related to Fig. 4A and 5) **(A)** The HMR C-terminus is required for HMR to form bright centromeric foci. Quantification of the percentage of centromeric foci (marked by CENP-C) associated with HMR in imunofluorescent stainings in SL2 cells expressing different *Hmr* transgenes (*Hmr* ^*+*^, *Hmr* ^*dC*^ and *Hmr* ^*2*^). Stainings were performed with DAPI, anti-HA (recognizing HA-HMR) and anti-CENP-C antibodies. For staining details refer to Fig.4A. **(B)** The number of centromeric foci per cell inversely correlates with HMR’s association with centromeres. Scatter plot displaying the relation between the percentage of centromeric foci associated with HMR (x-axis, binned by 10 % units) vs number of centromeric foci per cell (y-axis). Each dot represents a measured cell (a pool of all experiments from all Hmr alleles is displayed). **(C)** The ectopic expression of *Hmr* mutants correlates with higher numbers of centromeric foci. Boxplots displaying the number of centomeric foci per cell (y-axis) for each of the Hmr alleles (x-axis). Each dot represents a measured cell (a pool of all stained cells for each allele). **(D)** Protein expression in ovaries from *Hmr*^*ko*^ stocks complemented with different *Hmr* transgenes. Western blot probed with anti-HMR, anti-LHR and anti-LAMIN antibodies.

**Figure S7:**
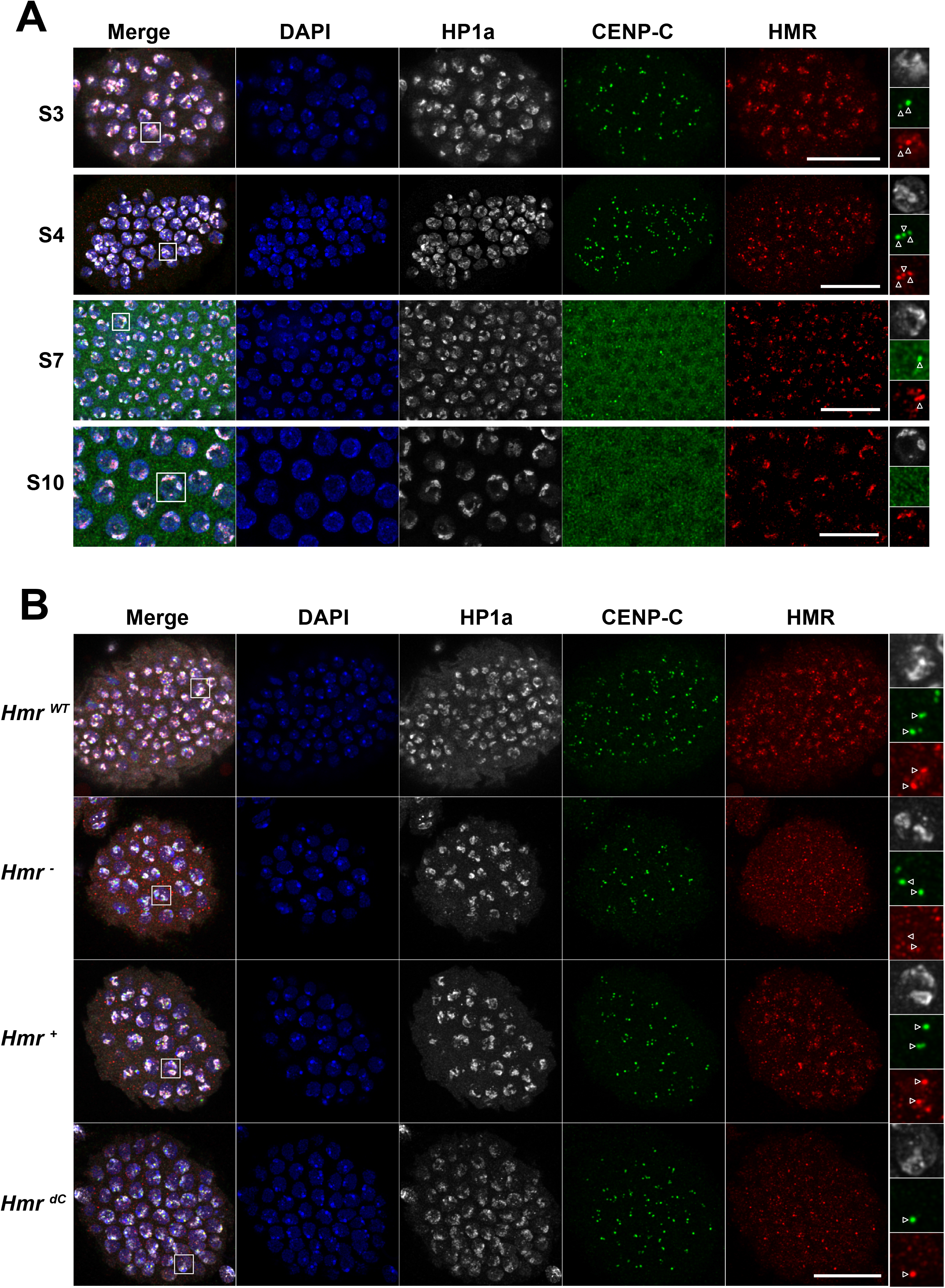
HMR localizes to the centromere in early-stage follicle cells and is mostly heterochromatic at later stages (related to Fig. 5C) **(A)** Localization of HMR in follicle cells isolated from egg chambers at different stages (**B)** Localization of different HMR variants in stage 4 follicle cells. Size bar indicates 15 µm.

## SUPPLEMENTARY METHODS

### Immunofluorescent staining in SL2 cells

For images in Fig. S2 leaky expression without copper induction was used in order to avoid overexpression artefacts. Cells were grown on #1.5H coverslips, fixed in PBS/ MeOH-free formaldehyde (4%) for 15 min at room temperature and permeabilized in PBS/Triton X-100 for 6 min on ice. After blocking with 5% normal goat serum dissolved in PBS (PBSN), coverslips were incubated with primary antibodies diluted in PBSN over night at 4°C. Fig. S2A: mouse anti-HA (Invitrogen 2-2.2.14; 1:300), rabbit anti-CID (Active Motif, 1:300), rat anti-HMR (2C10; 1:25); Fig. S2B: rat anti-HA (Sigma Aldrich 3F10; 1:100), mouse anti-HP1a (DSHB C1A9, 1:100), rabbit anti-CID (1:300). Following several washes with PBS/Triton X-100 0.05% and another blocking step with PBSN, secondary antibodies diluted in PBSN were added together with 1 µg/mL DAPI and incubated 1.5 hours at room temperature. Secondary antibodies: goat anti-rabbit Alexa Fluor Plus 647, donkey anti-rat Cy3, goat anti-mouse Alexa Fluor Plus 488. Following washes with PBS/Triton X-100 0.05% and PBS, coverslips were mounted in Prolong Diamond antifade.

### Immunofluorescent staining in ovaries

Fig. S7: All steps were performed as described in the main methods section except the following. Primary antibody solution (PBS-T, rat anti-HMR-2C10 1:20, mouse anti-HP1a-C1A9 1:10, rabbit anti-CENP-C 1:3000 and NDS 2%). Secondary antibody solution (200 µL PBS-T + donkey anti-mouse Alexa 488 1:600, donkey anti-rat Cy3 1:300, donkey anti-rabbit Alexa 647 and 2% NDS).

### Microscopy and downstream image analysis

Fig.4A: confocal microscopy z-scans were done on a Leica TCS SP5 (with 63x objective with 1.3 NA) with a step of 0.25 µM. Sum intensities projections were analyzed, and only cells with a minimum nucleoplasmic intensity of 70 a.u. on the anti-HMR channel were taken into account for analysis. Two different quantifications were performed. In one case cells were separated and counted based on the degree of co-localization between HMR and CENP-C: overlapping, partially overlapping or non-overlapping. In parallel, the number of CENP-C marked centromeric foci associated with HMR signal was measured. Both cells and centromeric foci were blind-counted, the experiment was repeated in 2 biological replicates and for each replicate at least two slides were measured (for each slide between 24 and 63 cells were quantified).

Confocal microscopy of data presented in Figures 5C and S2 was performed at the core facility bioimaging of the Biomedical Center using the following instruments and settings:

Figure S2: upright Leica SP8X WLL microscope (Klonike-upgraded) equipped with 405 nm laser, WLL2 laser (470 - 670 nm) and acusto-optical beam splitter. Images were acquired with a 63x/1.4 NA objective, pixel size set to 43 nm. The following spectral settings were used: DAPI (excitation 405 nm; emission 415 - 470 nm; detector: PMT), Alexa Fluor 488 (498 nm; 508 – 535 nm; HyD), Cy3 (551 nm; 561 – 610 nm; HyD), Alexa Fluor 647 (650 nm; 660 – 680 nm; HyD). Recording of image stacks was done at 200 Hz and line sequentially to avoid bleed-through. Hybrid detectors (HyDs) were operated in photon counting mode.

Figure 5C: inverted Leica SP8X STED 3D microscope (Klondike-upgraded), equipped with a 405 nm Laser, Argon Laser and a pulsed white light Laser (470 - 670 nm). Images were acquired with a 93x/1.3 NA Glycerol immersion objective, pixel size set to 63 nm. The following spectral settings were used: DAPI (405 nm; 415 - 470 nm; HyD), Alexa Fluor 488 (498 nm; 508 – 535 nm; HyD), Cy3 (551 nm; 560 – 604 nm; HyD), Alexa Fluor 647 (650 nm; 660 – 680 nm; HyD). Recording of image stacks was done at 200 Hz and line sequentially to avoid bleed-through. Hybrid detectors (HyDs) were operated in photon counting mode.

Image raw data was deconvolved using Huygens 17.10 p2. Further image processing, like generating maximum projections and linear adjustments of brightness and contrast, was done with ImageJ.

